# Reciprocal Disruptions in Thalamic and Hippocampal Resting-State Functional Connectivity in Youth with 22q11.2 Deletions

**DOI:** 10.1101/226951

**Authors:** Charles Schleifer, Amy Lin, Leila Kushan, Jie Lisa Ji, Genevieve Yang, Carrie E. Bearden, Alan Anticevic

## Abstract

22q11.2 deletion syndrome (22q11DS) is a recurrent copy number variant (CNV) with high penetrance for developmental neuropsychiatric disorders. Study of individuals with 22q11DS therefore may offer key insights into neural mechanisms underlying such complex illnesses. Resting-state functional MRI (rs-fMRI) studies in idiopathic schizophrenia have consistently revealed disruption of thalamic and hippocampal circuitry. Here, we sought to test whether this circuitry is similarly disrupted in the context of this genetic high-risk condition. To this end, resting-state functional connectivity patterns were assessed in a sample of young men and women with 22q11DS (n=42) and demographically matched healthy controls (n=39). Neuroimaging data were acquired via single-band protocols, and analyzed in line with methods provided by the Human Connectome Project (HCP). We computed functional relationships between individual-specific anatomically-defined thalamic and hippocampal seeds and all gray matter voxels in the brain. Whole-brain type I error protection was achieved through nonparametric permutation-based methods. 22q11DS patients displayed reciprocal disruptions in thalamic and hippocampal functional connectivity relative to control subjects. Thalamo-cortical coupling was increased in sensorimotor cortex, and reduced across associative networks. The opposite effect was observed for the hippocampus in regards to sensory and associative network connectivity. The thalamic and hippocampal dysconnectivity observed in 22q11DS suggest that high genetic risk for psychiatric illness is linked with disruptions in large-scale cortico-subcortical networks underlying higher-order cognitive functions. These effects highlight the translational importance of large-effect CNVs for informing mechanisms underlying neural disruptions observed in idiopathic developmental neuropsychiatric disorders.

**SIGNIFICANCE STATEMENT:** Investigation of neuroimaging biomarkers in highly penetrant genetic syndromes represents a more biologically tractable approach to identify neural circuit disruptions underlying developmental neuropsychiatric conditions. 22q11.2 deletion syndrome confers particularly high risk for psychotic disorders, and is thus an important translational model in which to investigate systems-level mechanisms implicated in idiopathic illness. Here, we show resting-state fMRI evidence of large-scale sensory and executive network disruptions in youth with 22q11DS. In particular, this study provides the first evidence that these networks are disrupted in a reciprocal fashion with regard to the functional connectivity of the thalamus and hippocampus, suggesting circuit-level dysfunction.

## INTRODUCTION

Remarkable genetic and clinical heterogeneity presents a challenge for mapping pathological processes underlying neuropsychiatric disorders such as schizophrenia and autism spectrum disorder (ASD). These disorders are increasingly viewed as developmental disruptions of neural circuitry with major genetic contributions (Insel, 2010; Geschwind and Flint, 2015). Thus, genetically-defined syndromes with strong predisposition for neuropsychiatric illness provide powerful models to elucidate neural mechanisms underlying these complex disorders.

22q11.2 Deletion Syndrome (22q11DS), also known as DiGeorge or Velocardiofacial syndrome (OMIM #188400, #192430), occurs in about 1 in 4,000 live births (McDonald-McGinn et al., 2015). It represents one of the greatest known genetic risk factors for psychosis, approximately 25 times population base rates (Bassett and Chow, 2008; Green et al., 2017), while additionally conferring elevated risk for multiple childhood disorders including attention deficit hyperactivity disorder, anxiety disorder, and ASD (Schneider et al., 2014).

Genes within the 22q11.2 locus are implicated in cortical circuit formation and functioning (Meechan et al., 2015; Paronett et al., 2015). Disrupted cortical interneuron migration has been observed in a 22q11.2 mouse model (Meechan et al., 2012; Toritsuka et al., 2013). Correspondingly, deletion carriers present with a range of structural and functional brain abnormalities, including cortical surface area reductions, altered white-matter microstructure (Kates et al., 2001; Jalbrzikowski et al., 2014; Schmitt et al., 2015), and, importantly, disruptions of large-scale network connectivity (Debbane et al., 2012; Padula et al., 2015). Recently, an independent components analysis approach revealed significant hypo-connectivity relative to controls within the anterior cingulate/precuneus and default mode networks, which reliably predicted 22q11DS case-control status in an independent cohort (Schreiner et al., 2017). Critically, due to its well-characterized genetic etiology, circuit-level abnormalities associated with 22q11.2 deletions can be experimentally manipulated in animals to generate causal links with circuit dysfunction. In humans, 22q11DS presents a compelling genetic high-risk model in which anomalous circuitry can be investigated prior to development of overt illness.

Specifically, aberrant connectivity of two key anatomically inter-connected structures, the thalamus and hippocampus, has been implicated in neuropsychiatric disorders (Brown et al., 2017) and schizophrenia in particular (Samudra et al., 2015). The thalamus serves as a critical hub for flow of sensory and higher-order information, facilitating information integration across cortical networks (Guo et al., 2017; Hwang et al., 2017). Consistent alterations of thalamo-cortical circuitry, involving a pattern of prefrontal-thalamic *hypo-connectivity,* concomitant with somatomotor-thalamic *hyper-connectivity,* have been identified in schizophrenia patients and at-risk youth (Welsh et al., 2010; Woodward et al., 2012; Anticevic et al., 2014). Similarly, the hippocampus features prominently in schizophrenia neurobiology (Weinberger, 1987). Post-mortem schizophrenia studies have demonstrated hippocampal alterations in excitatory pyramidal cells and local inhibitory interneurons. Hippocampal-prefrontal dysconnectivity during cognitive processing has been proposed as a translational phenotype for schizophrenia, as evidenced by a 22q11 mouse model (Mukai et al., 2015) and by findings of altered connectivity in those at familial high risk for the illness (Meyer-Lindenberg, 2010). Critically, the thalamus and hippocampus exhibit opposing resting-state connectivity patterns in healthy adults (Stein et al., 2000), which would predict distinct alterations in a genetic risk model based on a CNV that disrupts neural circuits. Yet, these translational neural phenotypes have not been investigated in a genetic risk model such as 22q11DS.

Here we take a hypothesis-based approach to study large-scale network alterations in 22q11DS by leveraging findings from animal models of the disorder and imaging work in humans. Using the Human Connectome Project analytical pipeline, which yields exceptional cortical spatial alignment (Glasser et al., 2013), we computed functional relationships between subject-specific anatomically-defined thalamic and hippocampal seeds in 22q11DS youth and matched controls. Relative to controls, 22q11DS youth exhibited thalamo-cortical *hyper-connectivity* with sensorimotor cortex but *hypo-connectivity* with associative networks. An opposing (i.e. interactive) pattern was found for hippocampal-cortical circuitry, supporting a 22q11DS neural phenotype with distinct effects on thalamic and hippocampal circuits.

## METHODS

### Participants

The total sample consisted of 81 participants (7 to 26 years of age; 42 22q11DS and 39 demographically matched healthy controls), recruited from an ongoing longitudinal study at the University of California, Los Angeles (UCLA). 22q11DS participants all had a molecularly confirmed 22q11.2 deletion (see **Table 1** for demographic details). Exclusion criteria for all study participants were: neurological or medical condition disorder that might affect performance, insufficient fluency in English, and/or substance or alcohol abuse and/or dependence within the past 6 months. Healthy controls (HCS) additionally could not meet diagnostic criteria for any major mental disorder, based on information gathered during administration of the Structured Clinical Interview for DSM-IV Axis I Disorders (First et al., 1996). After study procedures had been fully explained, adult participants provided written consent, while participants under the age of 18 years provided written assent with the written consent of their parent or guardian. The UCLA Institutional Review Board (IRB) approved all study procedures and informed consent documents.

**Table 1.**
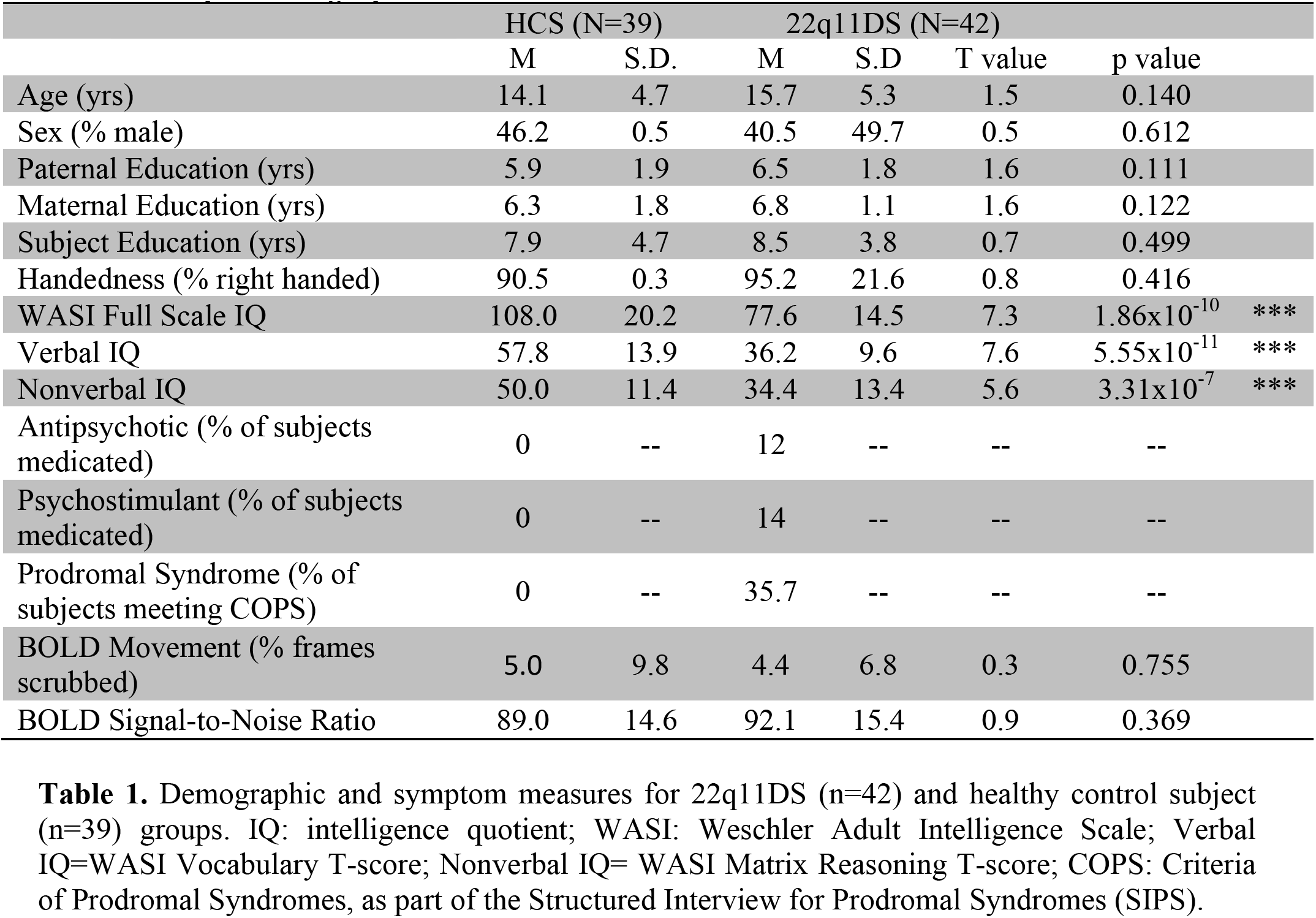
Demographic and symptom measures for 22q11DS (n=42) and healthy control subject (n=39) groups. IQ: intelligence quotient; WASI: Weschler Adult Intelligence Scale; Verbal IQ=WASI Vocabulary T-score; Nonverbal IQ= WASI Matrix Reasoning T-score; COPS: Criteria of Prodromal Syndromes, as part of the Structured Interview for Prodromal Syndromes (SIPS).

### Neuroimaging Acquisition

All subjects were imaged on a 3-Tesla Siemens TimTrio scanner with a 32-channel phased array head coil at the UCLA Center for Cognitive Neuroscience (CCN). Resting BOLD images were acquired in 34 interleaved axial slices parallel to the anterior-posterior commissure (AC-PC) using a fast gradient-echo, echo-planar sequence [voxel size=3x3x4mm, time repetition (TR) = 2000ms, time echo (TE) = 30ms, flip angle = 90°, matrix = 64x64, field of view = 192x192 mm]. Acquisition lasted 5.1 minutes and produced 152 volumes. High-resolution T1w images were collected in 160 sagittal slices via a magnetization-prepared rapid gradient-echo sequence (MP-RAGE) [voxel size = 1x1x1mm, TR = 2300ms, TE = 2.91ms, flip angle = 9°, matrix = 240 x 256, field of view = 240x256 mm].

### Clinical Assessment

On the same day as the scan, demographic information and clinical measures were collected for each participant by trained master’s level clinicians (see **Table 1**). Verbal IQ was assessed via the Wechsler Abbreviated Scale of Intelligence (WASI) Vocabulary subtest and Non-verbal IQ was assessed via the WASI Matrix Reasoning subtest. Psychiatric and dimensional psychotic-like symptoms were assessed via the Structured Interview for Prodromal Symptoms (SIPS; Tandy J. Miller et al., 2002). See (Jalbrzikowski et al., 2012; Jalbrzikowski et al., 2013) for more details on study ascertainment and recruitment procedures.

### Data Preprocessing

Structural and functional MRI data were first preprocessed according the methods provided by the Human Connectome Project (HCP), outlined below, and described in detail by the WU-Minn HCP consortium (Glasser et al., 2013). These open-source HCP algorithms, which we further optimized for compatibility with legacy single-band data in this study, represent the current state-of-the-art approaches in spatial distortion correction, registration, and maximization of high-resolution signal-to-noise (SNR) (Glasser et al., 2016). All processing methods closely followed the minimal processing pipelines as outlined by Glasser and colleagues (Glasser et al., 2013), with a few key modifications.

The adapted HCP pipeline included the following steps: i) the T1-weighted images were corrected for bias-field distortions and warped to the standard Montreal Neurological Institute-152 (MNI-152) brain template through a combination of linear and non-linear transformations using the FMRIB Software Library (FSL) linear image registration tool (FLIRT) and non-linear image registration tool (FNIRT) (Jenkinson et al., 2002). ii) FreeSurfer’s recon-all pipeline was employed to compute brain-wide segmentation of gray and white matter to produce individual cortical and subcortical anatomical segmentation (Reuter et al., 2012). iii) Next, cortical surface models were generated for pial and white matter boundaries as well as segmentation masks for each subcortical grey matter voxel. Using the pial and white matter surface boundaries, a ‘cortical ribbon’ was defined along with corresponding subcortical voxels, which were combined to generate the Connectivity Informatics Technology Initiative (CIFTI) volume/surface ‘gray-ordinate’ space for each individual subject, which drastically reduces file management for combined surface and volume analyses and visualization and establishes a combined cortical surface and subcortical volume coordinate system (Glasser et al., 2013). iv) the cortical surfaces were then registered to the group average HCP atlas using surface-based registration based on cortical landmark features, whereas the subcortical ‘volume’ component of the image was brought into group atlas alignment via non-linear registration (Glasser et al., 2013). v) The BOLD data were motion corrected and aligned to the middle frame of every run via FLIRT. In turn, a liberal brain-mask was applied to exclude signal from non-brain tissue. After initial processing in NIFTI volume space, BOLD data were converted to the CIFTI gray matter matrix by sampling from the anatomically-defined gray matter cortical ribbon whereas the subcortical voxels were isolated using subject-specific FreeSurfer segmentation. The subcortical volume component of the BOLD data was then aligned to the group atlas as part of the NIFTI processing in a single transform step that concatenates all of the transform matrixes for each prior processing step (i.e. motion correction, registration, distortion correction). This produced a single nonlinear transformation to minimize interpolation cost. In turn, the cortical surface component of the CIFTI file was aligned to the HCP atlas using surface-based nonlinear deformation based on sulcal features.

Following these ‘minimal’ HCP preprocessing steps, a high-pass filter (>0.5 Hz) was applied to the BOLD time series in order to remove low temporal frequencies and scanner drift. In-house MATLAB tools were then used to compute the signal in the ventricles, deep white matter, and across all gray matter voxels (proxy of global mean signal regression to address spatially pervasive sources of artifacts; (Power et al., 2017)). These time series were modeled as nuisance variables and were regressed out of the gray matter voxels. Subsequent analyses used the residual BOLD time series following these de-noising steps.

### Functional Connectivity Analyses

Thalamic and hippocampal seeds were first defined individually for each subject through automatic anatomical segmentation of high-resolution structural images via FreeSurfer software as part of the HCP minimal preprocessing pipelines. These structures were then used as ‘seeds’, as conducted in our prior work (Anticevic et al., 2014). Specifically, Pearson correlations were computed between the mean BOLD signal time series in each seed and the BOLD time series at every other cortical and subcortical vertex in CIFTI gray-ordinate space. These correlation maps were then standardized for statistical analyses via Fisher r-to-Z transformation.

As noted, the thalamus and hippocampus in humans exhibit distinct resting-state connectivity profiles (Stein et al., 2000). In fact, this would predict distinct alterations in a genetic risk model based on a de novo CNV that uniformly affects neural circuits. In turn, combining two ‘seed’ regions, both of which may be affected, but with opposing predicted directions of alterations, constitutes a more powered neural marker. Put differently, we hypothesized a *Group* by *Seed* interaction whereby 22q11DS may exhibit distinct bi-directional alterations across the hippocampal and thalamic systems. To confirm the viability of this logic, we computed an *a priori* quantitative independent test of differences in thalamic and hippocampal connectivity in a sample of 339 unrelated healthy adults derived from the Human Connectome Project (HCP). This provided the basis for the expected interactive effects between thalamic and hippocampal seeds in the core between-group analysis. In other words, the purpose of the HCP dataset here was to serve as a large normative sample to provide an empirical independent basis for the proposed clinical hypotheses.

Next, to test the *Group x Seed* interaction effect with the BOLD rs-fcMRI as the dependent measure, we computed a two-way repeated measures ANOVA with a factor of *Group* (22q11DS vs. HCS) and *Seed* (thalamus vs. hippocampus). Whole-brain type I error protection was applied via non-parametric permutation testing with FSL’s Permutation Analysis of Linear Models (PALM) algorithm (Winkler et al., 2014) with 10,000 permutations. This approach circumvents the distributional assumptions (e.g. normality) that may result in type I error inflation (Eklund et al., 2016). We also independently repeated our seed-based analyses in each of seven *a priori* functional networks described by Yeo and colleagues to test for network specificity of the hypothesized effects (Buckner et al., 2011; Yeo et al., 2011; Choi et al., 2012).

To quantify the differential contributions of thalamic and hippocampal sub-regions, a k-means algorithm was used to cluster voxels within each seed based on the correlation distance between their group-level rs-fcMRI effects. Multiple cluster solutions were possible, but we elected to focus on a parsimonious two-cluster solutions for the thalamus and hippocampus because the higher cluster solutions explained proportionally less of the variance, as demonstrated by the ‘Elbow Method’ (Thorndike, 1953). Seed-based functional connectivity was subsequently computed for each of the four resultant clusters (two per seed). Each cluster’s whole-brain connectivity matrix was then correlated with the whole-brain connectivity matrix previously computed for the whole seed. Within the thalamus and hippocampus, the two resulting Pearson coefficients were compared using Steiger’s Z-test to determine which cluster’s connectivity profile most resembled that of the whole seed (Steiger, 1980). For the thalamus, the connectivity profiles of the whole seed, as well as the anterior and posterior data-derived clusters, were compared to the functional connectivity of seven *a priori* anatomical seeds derived from the FSL thalamic atlas.

The utility of the observed rs-fcMRI effects for individual classification accuracy was assessed via a supervised binary classification algorithm. A total n=1000 iterations of a Support Vector Machine (SVM) were computed, each randomly splitting the n=81 pooled subjects and training on n=41, then using split-half crossvalidation with the remaining n=40 to build a distribution of receiver operator (ROC) curves. One-dimensional SVMs were trained and tested on a single factor consisting of the linear combination of thalamic and hippocampal connectivity to each of the interaction-derived ROIs ([thalROIa + hippROIb] - [thalROIb + hippROIa]). This was repeated for the network-derived results ([thalSOM + hippFPN] - [thalFPN + hippSOM]).

Several analyses were performed in order to address confounds potentially introduced during data acquisition or processing. For BOLD images, frames with significant head movement were flagged based on algorithms and intensity thresholds recommended for multi-band data (Power et al., 2012). Temporal signal-to-noise ratio (SNR) was calculated for each subject as the ratio of mean BOLD signal to its standard deviation over time. Movement (percentage of flagged frames) and SNR correlations with rs-fcMRI effects were examined for both groups. To assess medication as a potential confound, two-sample t-tests were computed between rs-fcMRI effects in medicated versus unmediated 22q11DS patients for the subsets taking antipsychotic medications and dopaminergic stimulants. Global signal regression (GSR) was included as a preprocessing step for the main analyses, but functional connectivity was also re-computed for the data without GSR in order to ensure that the effects were comparable at the whole-brain level, and within the specific ROIs derived from permutation testing.

## RESULTS

### 22q11DS Is Associated with Distinct Functional Dysconnectivity for Thalamus and Hippocampus

As noted, we sought to test if 22q11DS is characterized by disruptions in thalamic as well as hippocampal resting-state functional connectivity (rs-fcMRI). We hypothesized dissociable effects across thalamic and hippocampal seeds, given their known differences in functional connectivity patterns. To establish this effect, we first conducted a ‘control’ analysis in the n=339 healthy adult subjects collected by the HCP (**Figure 1**). Results showed that the rs-fcMRI profiles of the thalamus and hippocampus are intrinsically anti-correlated with respect to a broad set of regions overlapping with sensory and executive networks.

**Figure 1.**
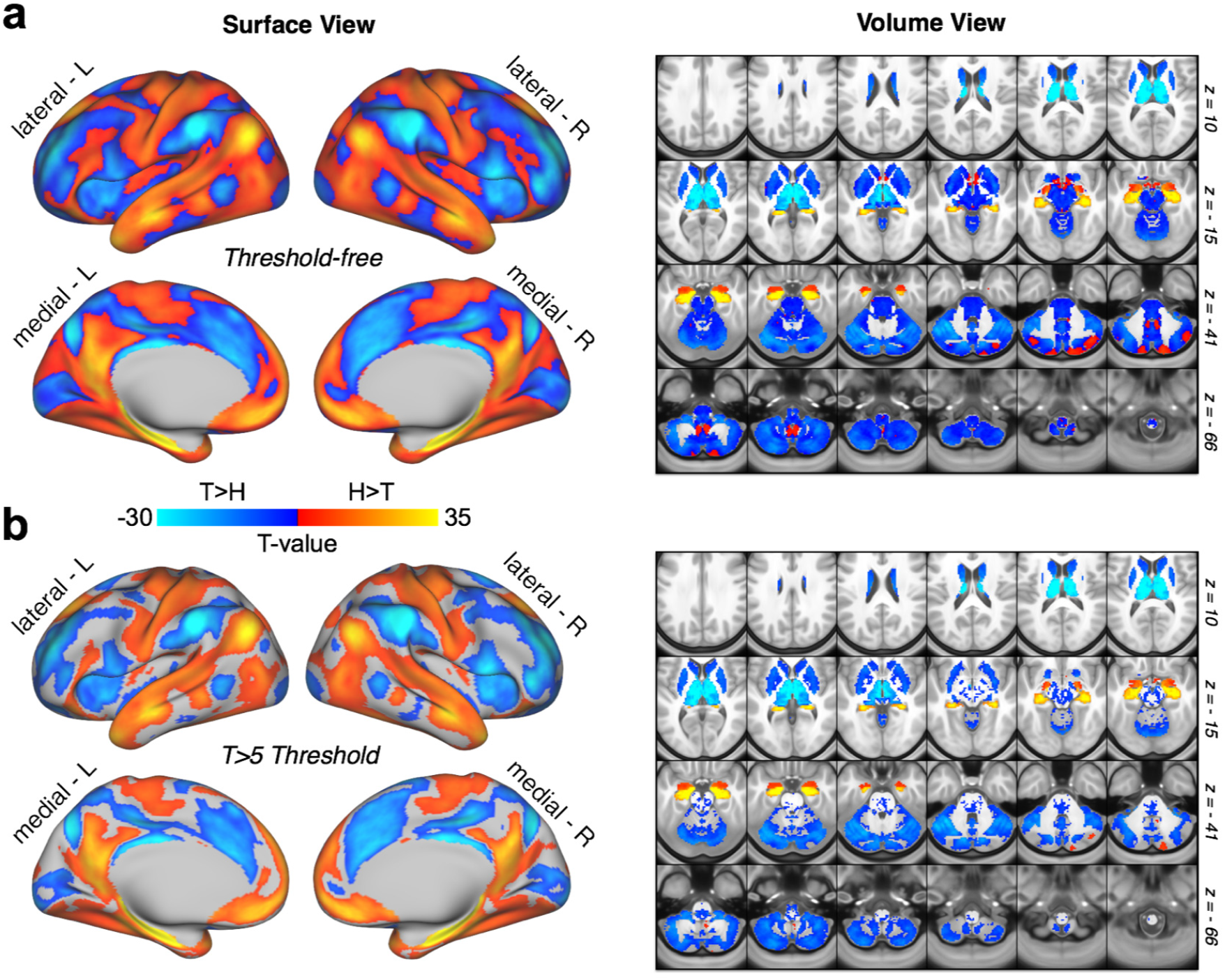
Hippocampal vs. Thalamic Seed Connectivity, N=339 Healthy Adults (HCP Dataset). Comparison of thalamic and hippocampal resting-state functional connectivity in the Human Connectome Project dataset. **(a)** Surface and volume maps showing the threshold-free dependent-samples t-test between thalamic and hippocampal functional connectivity in n=339 healthy adult subjects. **(b)** The same contrast, masked at T value > 5.

Next, we tested whether these rs-fcMRI disruptions exhibit interactive effects for 22q11DS versus HCS. As predicted, there were two sets of regions exhibiting a significant 2x2 *Group* by *Seed* interaction: i) sensory-motor regions, marked by hyper-connectivity for the thalamus but hypo-connectivity with the hippocampus; ii) a cerebellar region marked by hypo-connectivity for the thalamus but hyper-connectivity with the hippocampus (**Figure 2**; see **Table 4** for all regions surviving type I error correction). Put differently, the interaction was driven by the 22q11DS group exhibiting significantly increased thalamic connectivity (but decreased hippocampal connectivity) with bilateral sensorimotor regions, including the pre- and postcentral gyri and superior temporal gyrus, whereas the opposite effect (decreased thalamic and increased hippocampal connectivity) was observed for a region in the left cerebellum. While this effect was localized to the cerebellum following type I error correction, the threshold-free maps show a broader set of prefrontal and parietal regions that trend towards significance (**Figure 3**).

**Figure 2.**
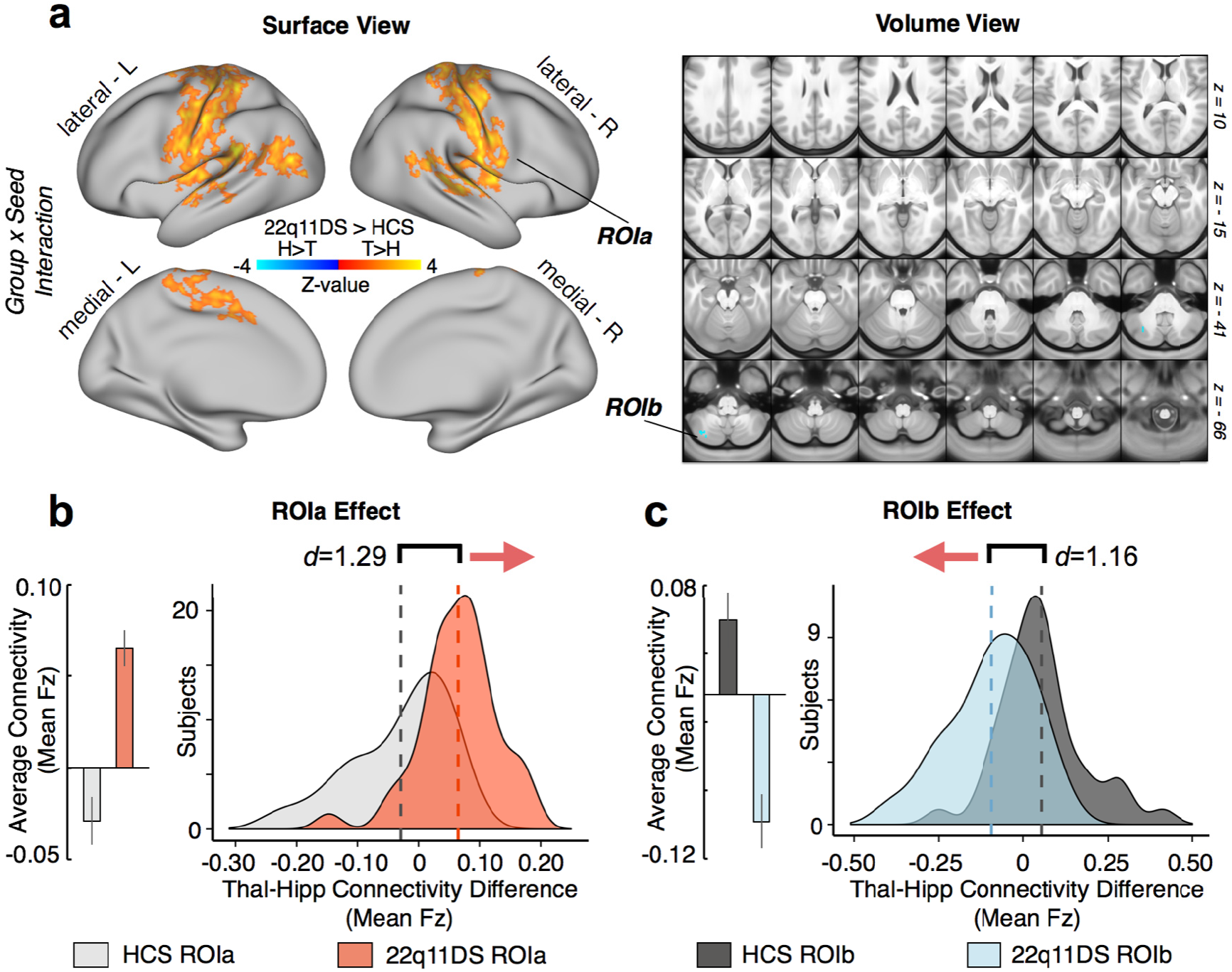
Interaction-Derived Effects. Resting-state functional connectivity of the thalamus and hippocampus in 22q11DS and healthy controls (HCS). **(a)** Surface and volume maps showing type I error-protected group-level contrast for the 2x2 interaction between *Group* (22q11DS vs. HCS) and *Seed* (thalamus vs. hippocampus). Yellow-orange (ROIa) indicates an effect whereby 22q11DS showed thalamic hyper-connectivity but hippocampal hypo-connectivity relative to HCS. The blue contrast (ROIb) indicates an effect whereby 22q11DS showed thalamic hypo-connectivity but hippocampal hyper-connectivity relative to HCS. **(b)** Difference scores between thalamic and hippocampal connectivity to ROIa (mean Fz) across subjects in each group. Group means (left) and distributions (right) shown to illustrate the direction of the effect. **(c)** Same as **(b)** for ROIb, showing an effect in the opposite direction as ROIa. Note: histograms are based on the data extracted from the maps presented in **(a)**. Individual thalamic and hippocampal effects are presented in Figure 10 in comparison with functional network-derived effects.

**Figure 3.**
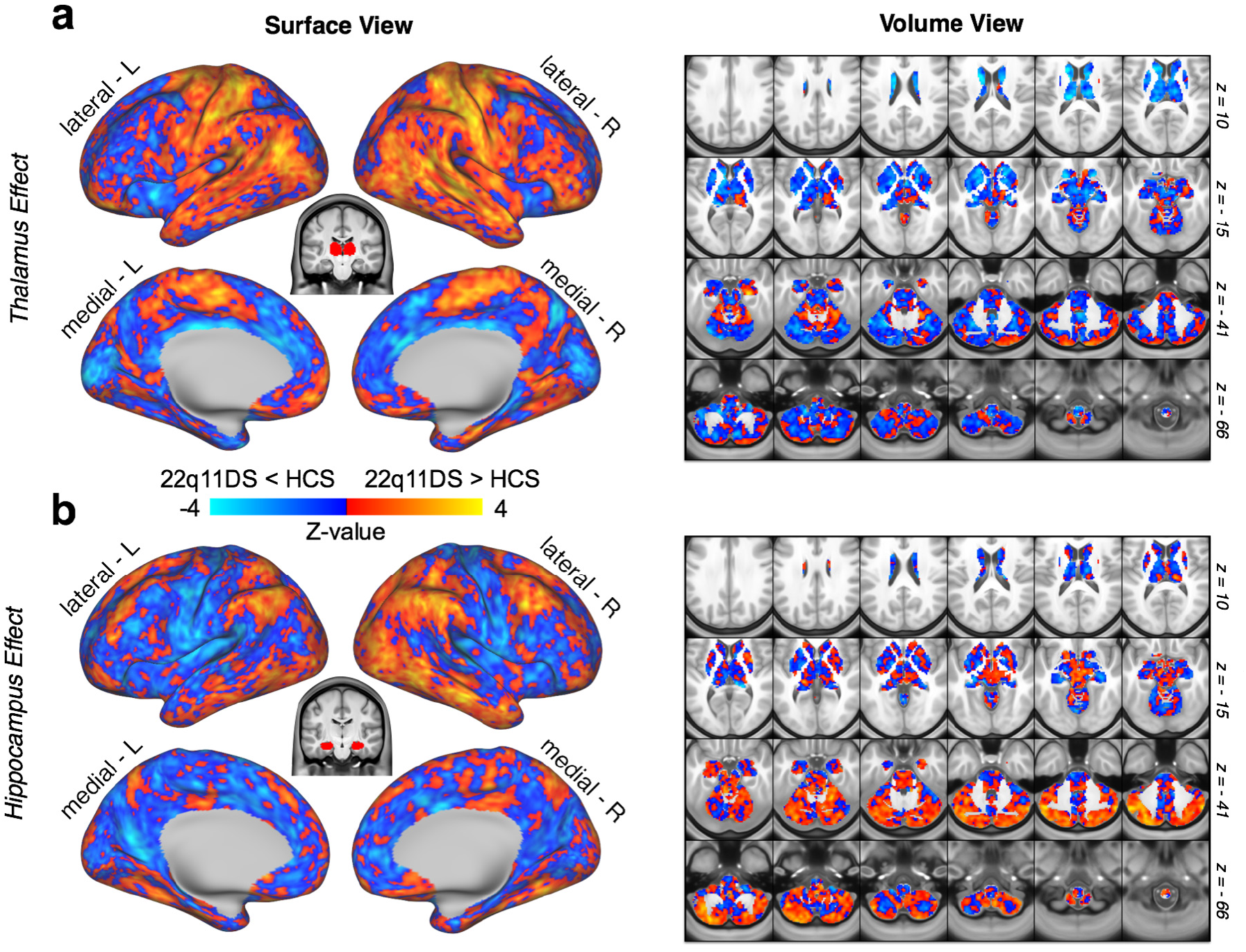
Threshold-Free Seed-Based Connectivity Maps. Threshold-free group contrasts for thalamic and hippocampal functional connectivity. **(a)** Surface and volume maps showing the threshold-free effect (22q11DS vs. HCS) for the thalamic seed. The yellow-orange contrast indicates 22q11DS > HCS. Blue indicates HCS > 22q11DS. **(b)** Same effect shown for the hippocampal seed.

### Ruling out Motion, SNR, and Medication Effects

To ensure that the observed effects were not attributable to differential motion between groups, or to differential SNR profiles, we correlated both measures with the functional connectivity values for both seeds to both interaction-derived ROIs, as well as with the linear combination of these four connectivity values. No significant relationships were observed between functional connectivity and motion or SNR for the 22q11DS or HCS groups (see **Table 2**). Within the 22q11DS group, mean rs-fcMRI effects were also compared between cohorts of medicated and un-medicated patients (with regards to antipsychotic and stimulant medication). No significant effects of either medication were observed (see **Table 3**).

**Table 2.**
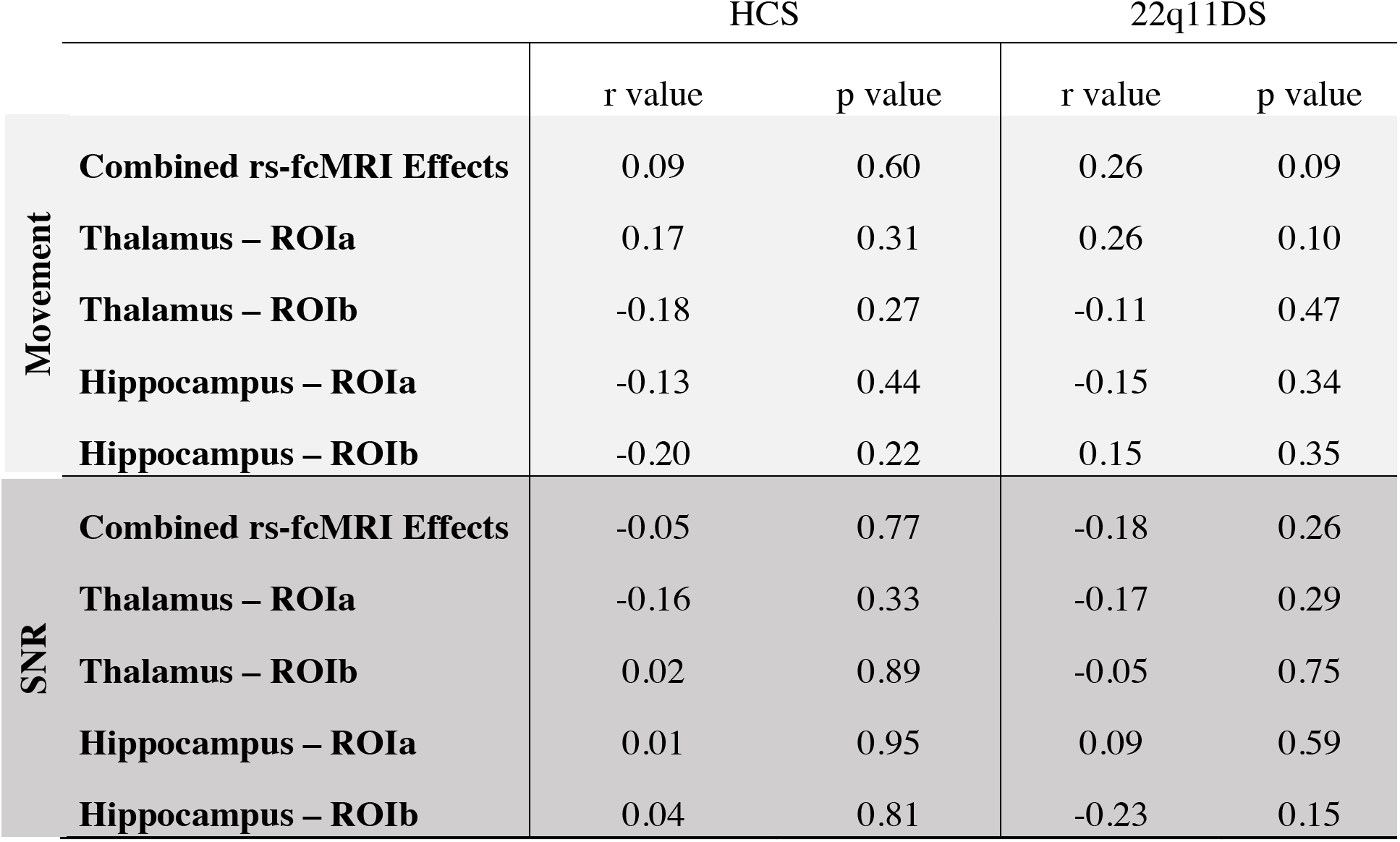
Movement and SNR Relationships. Pearson correlations showing no significant relationship between rs-fcMRI effects (mean Fz connectivity values) and measures of head movement signal-to-noise ratio (SNR). ‘Combined fcMRI Effects’ refers to the linear combination of connectivity values from the thalamus and hippocampus to ROIa and ROIb ([thalROIa + hippROIb] - [thalROIb + hippROIa]). Connectivity between each seed and ROI (e.g. Thalamus to ROIa) was also individually tested for correlation with motion and SNR.

**Table 3.** Comparison of fcMRI effects in medicated versus un-medicated 22q11DS subjects, with regards to antipsychotics and dopaminergic stimulants. Two-sample t-tests are shown for the linear combination of connectivity values from both seeds and ROIs ([thalROIa + hippROIb] - [thalROIb + hippROIa]), as well as for the connectivity of each individual seed to each ROI. No significant effects of medication were observed.

|  | Antipsychotic |  | Stimulant |  |
| --- | --- | --- | --- | --- |
|  | T-Value | p-Value | T-Value | p-Value |
| **Combined fcMRI Effects** | -1.643 | 0.144 | 0.197 | 0.849 |
| **Thalamus – RO Ia** | -0.342 | 0.735 | -0.581 | 0.581 |
| **Thalamus – RO Ib** | 1.003 | 0.361 | 0.390 | 0.710 |
| **Hippocampus – RO Ia** | -0.560 | 0.603 | -1.002 | 0.346 |
| **Hippocampus – RO Ib** | -1.131 | 0.302 | 0.609 | 0.565 |

**Table 4.** Pairwise Comparisons - Region Coordinates, P-values & Effect Sizes (Interaction-Derived Effect)

| X | Y | Z | Hemisphere | Landmark | Size | Comparison | *d* | Mean *T* | *p* | Comparison | *d* | Mean *T* | *p* |
| --- | --- | --- | --- | --- | --- | --- | --- | --- | --- | --- | --- | --- | --- |
| -5 | 0 | 43 | cortex left | cingulate gyrus | 348 mm2 | Pt-Con Thal. | 0.24 | -1.09 | 0.279 | Pt-Con Hipp. | 0.58 | 2.62 | 0.010 |
| -13 | -37 | 75 | cortex left | postcentral gyrus | 376 mm2 | Pt-Con Thal. | 0.45 | -2.01 | 0.048 | Pt-Con Hipp. | 0.40 | 1.82 | 0.073 |
| -8 | -35 | 60 | cortex left | paracentral lobule | 268 mm2 | Pt-Con Thal. | 0.58 | -2.59 | 0.011 | Pt-Con Hipp. | 0.27 | 1.22 | 0.225 |
| -9 | -26 | 51 | cortex left | cingulate sulcus | 304 mm2 | Pt-Con Thal. | 0.36 | -1.61 | 0.111 | Pt-Con Hipp. | 0.48 | 2.14 | 0.036 |
| -6 | -26 | 43 | cortex left | paracentral lobule | 268 mm2 | Pt-Con Thal. | 0.51 | -2.29 | 0.025 | Pt-Con Hipp. | 0.18 | 0.80 | 0.424 |
| -6 | -4 | 56 | cortex left | paracentral lobule | 372 mm2 | Pt-Con Thal. | 0.60 | -2.69 | 0.009 | Pt-Con Hipp. | 0.29 | 1.32 | 0.189 |
| -37 | -7 | 65 | cortex left | precentral sulcus | 220 mm2 | Pt-Con Thal. | 0.50 | -2.23 | 0.029 | Pt-Con Hipp. | 0.32 | 1.44 | 0.154 |
| -18 | -19 | 76 | cortex left | precentral gyrus | 420 mm2 | Pt-Con Thal. | 0.41 | -1.84 | 0.069 | Pt-Con Hipp. | 0.42 | 1.89 | 0.063 |
| -16 | -5 | 74 | cortex left | superior frontal gyrus | 404 mm2 | Pt-Con Thal. | 0.39 | -1.77 | 0.081 | Pt-Con Hipp. | 0.51 | 2.29 | 0.025 |
| -58 | -18 | 46 | cortex left | postcentral gyrus | 684 mm2 | Pt-Con Thal. | 0.72 | -3.23 | 0.002 | Pt-Con Hipp. | 0.50 | 2.26 | 0.026 |
| -39 | -43 | 62 | cortex left | superior parietal lobule | 944 mm2 | Pt-Con Thal. | 0.92 | -4.15 | 0.000 | Pt-Con Hipp. | 0.17 | 0.78 | 0.436 |
| -29 | -35 | 72 | cortex left | postcentral gyrus | 336 mm2 | Pt-Con Thal. | 0.53 | -2.38 | 0.020 | Pt-Con Hipp. | 0.32 | 1.46 | 0.148 |
| -39 | -26 | 64 | cortex left | central sulcus | 592 mm2 | Pt-Con Thal. | 0.52 | -2.35 | 0.021 | Pt-Con Hipp. | 0.60 | 2.72 | 0.008 |
| -49 | -9 | 49 | cortex left | central sulcus | 556 mm2 | Pt-Con Thal. | 0.33 | -1.49 | 0.140 | Pt-Con Hipp. | 0.73 | 3.27 | 0.002 |
| -64 | -28 | 0 | cortex left | superior temporal gyrus | 200 mm2 | Pt-Con Thal. | 0.33 | -1.48 | 0.143 | Pt-Con Hipp. | 0.59 | 2.64 | 0.010 |
| -55 | -39 | 22 | cortex left | posterior sylvian fissure | 824 mm2 | Pt-Con Thal. | 0.61 | -2.76 | 0.007 | Pt-Con Hipp. | 0.76 | 3.43 | 0.001 |
| -61 | -26 | 6 | cortex left | medial temporal lobe | 408 mm2 | Pt-Con Thal. | 0.41 | -1.84 | 0.069 | Pt-Con Hipp. | 0.77 | 3.46 | 0.001 |
| -42 | -12 | 8 | cortex left | insula | 560 mm2 | Pt-Con Thal. | 0.20 | -0.91 | 0.367 | Pt-Con Hipp. | 0.97 | 4.34 | 0.000 |
| -46 | -86 | 3 | cortex left | inferior occipital gyrus | 92 mm2 | Pt-Con Thal. | 0.63 | -2.82 | 0.006 | Pt-Con Hipp. | 0.21 | 0.95 | 0.348 |
| -49 | -77 | 13 | cortex left | superior occipital gyrus | 696 mm2 | Pt-Con Thal. | 0.73 | -3.29 | 0.001 | Pt-Con Hipp. | 0.27 | 1.24 | 0.220 |
| -53 | -17 | 16 | cortex left | sylvian fissure | 728 mm2 | Pt-Con Thal. | 0.26 | -1.16 | 0.250 | Pt-Con Hipp. | 0.81 | 3.63 | 0.001 |
| -62 | -30 | 40 | cortex left | supramarginal gyrus | 348 mm2 | Pt-Con Thal. | 0.54 | -2.44 | 0.017 | Pt-Con Hipp. | 0.27 | 1.24 | 0.220 |
| -66 | -10 | 25 | cortex left | postcentral gyrus | 468 mm2 | Pt-Con Thal. | 0.40 | -1.78 | 0.079 | Pt-Con Hipp. | 0.73 | 3.27 | 0.002 |
| -60 | -7 | 33 | cortex left | central sulcus | 536 mm2 | Pt-Con Thal. | 0.39 | -1.75 | 0.084 | Pt-Con Hipp. | 0.63 | 2.81 | 0.006 |
| -56 | 5 | 5 | cortex left | sylvian fissure | 372 mm2 | Pt-Con Thal. | 0.31 | -1.40 | 0.165 | Pt-Con Hipp. | 0.80 | 3.61 | 0.001 |
| -52 | -10 | -5 | cortex left | medial temporal lobe | 256 mm2 | Pt-Con Thal. | 0.43 | -1.93 | 0.057 | Pt-Con Hipp. | 0.72 | 3.26 | 0.002 |
| -53 | 0 | -12 | cortex left | medial temporal lobe | 172 mm2 | Pt-Con Thal. | 0.47 | -2.10 | 0.039 | Pt-Con Hipp. | 0.54 | 2.41 | 0.018 |
| -61 | -10 | -2 | cortex left | superior temporal gyrus | 284 mm2 | Pt-Con Thal. | 0.35 | -1.58 | 0.119 | Pt-Con Hipp. | 0.73 | 3.26 | 0.002 |
| -62 | -32 | -9 | cortex left | superior temporal sulcus | 208 mm2 | Pt-Con Thal. | 0.46 | -2.06 | 0.043 | Pt-Con Hipp. | 0.26 | 1.15 | 0.253 |
| 9 | -25 | 72 | cortex right | paracentral lobule | 144 mm2 | Pt-Con Thal. | 0.46 | -2.08 | 0.041 | Pt-Con Hipp. | 0.28 | 1.25 | 0.215 |
| 45 | -10 | 59 | cortex right | precentral gyrus | 352 mm2 | Pt-Con Thal. | 0.65 | -2.91 | 0.005 | Pt-Con Hipp. | 0.49 | 2.22 | 0.029 |
| 18 | -34 | 76 | cortex right | postcentral gyrus | 260 mm2 | Pt-Con Thal. | 0.48 | -2.15 | 0.035 | Pt-Con Hipp. | 0.42 | 1.89 | 0.062 |
| 31 | -13 | 72 | cortex right | precentral gyrus | 368 mm2 | Pt-Con Thal. | 0.84 | -3.78 | 0.000 | Pt-Con Hipp. | 0.19 | 0.84 | 0.401 |
| 15 | -21 | 76 | cortex right | precentral gyrus | 184 mm2 | Pt-Con Thal. | 0.31 | -1.41 | 0.163 | Pt-Con Hipp. | 0.55 | 2.49 | 0.015 |
| 55 | -3 | 46 | cortex right | central sulcus | 164 mm2 | Pt-Con Thal. | 0.20 | -0.88 | 0.380 | Pt-Con Hipp. | 0.52 | 2.32 | 0.023 |
| 29 | -29 | 70 | cortex right | central sulcus | 440 mm2 | Pt-Con Thal. | 0.39 | -1.75 | 0.083 | Pt-Con Hipp. | 0.74 | 3.32 | 0.001 |
| 48 | -27 | 64 | cortex right | postcentral gyrus | 444 mm2 | Pt-Con Thal. | 0.75 | -3.37 | 0.001 | Pt-Con Hipp. | 0.18 | 0.81 | 0.423 |
| 45 | -21 | 59 | cortex right | central sulcus | 624 mm2 | Pt-Con Thal. | 0.59 | -2.68 | 0.009 | Pt-Con Hipp. | 0.62 | 2.78 | 0.007 |
| 59 | -45 | 14 | cortex right | superior temporal sulcus | 468 mm2 | Pt-Con Thal. | 0.83 | -3.73 | 0.000 | Pt-Con Hipp. | 0.26 | 1.18 | 0.241 |
| 65 | -34 | 6 | cortex right | superior temporal gyrus | 388 mm2 | Pt-Con Thal. | 0.81 | -3.62 | 0.001 | Pt-Con Hipp. | 0.46 | 2.05 | 0.044 |
| 50 | -27 | 11 | cortex right | posterior sylvian fissure | 336 mm2 | Pt-Con Thal. | 0.12 | -0.55 | 0.581 | Pt-Con Hipp. | 0.89 | 4.00 | 0.000 |
| 44 | -1 | 3 | cortex right | insula | 380 mm2 | Pt-Con Thal. | 0.48 | -2.16 | 0.034 | Pt-Con Hipp. | 0.58 | 2.60 | 0.011 |
| 20 | -44 | 75 | cortex right | postcentral sulcus | 444 mm2 | Pt-Con Thal. | 0.56 | -2.50 | 0.014 | Pt-Con Hipp. | 0.45 | 2.04 | 0.044 |
| 31 | -50 | 70 | cortex right | superior parietal lobule | 360 mm2 | Pt-Con Thal. | 0.78 | -3.49 | 0.001 | Pt-Con Hipp. | 0.12 | 0.54 | 0.589 |
| 58 | -59 | 16 | cortex right | angular gyrus | 300 mm2 | Pt-Con Thal. | 0.60 | -2.70 | 0.009 | Pt-Con Hipp. | 0.39 | 1.75 | 0.084 |
| 68 | -9 | 21 | cortex right | postcentral gyrus | 488 mm2 | Pt-Con Thal. | 0.63 | -2.84 | 0.006 | Pt-Con Hipp. | 0.54 | 2.43 | 0.018 |
| 59 | -12 | 47 | cortex right | precentral gyrus | 564 mm2 | Pt-Con Thal. | 0.63 | -2.83 | 0.006 | Pt-Con Hipp. | 0.42 | 1.88 | 0.064 |
| 60 | -4 | 34 | cortex right | central sulcus | 588 mm2 | Pt-Con Thal. | 0.49 | -2.20 | 0.031 | Pt-Con Hipp. | 0.56 | 2.51 | 0.014 |
| 56 | -7 | 13 | cortex right | sylvian fissure | 460 mm2 | Pt-Con Thal. | 0.41 | -1.82 | 0.072 | Pt-Con Hipp. | 0.64 | 2.86 | 0.005 |
| 55 | 4 | -14 | cortex right | medial temporal lobe | 196 mm2 | Pt-Con Thal. | 0.61 | -2.75 | 0.007 | Pt-Con Hipp. | 0.57 | 2.55 | 0.013 |
| 50 | -17 | 3 | cortex right | medial temporal lobe | 196 mm2 | Pt-Con Thal. | 0.49 | -2.19 | 0.031 | Pt-Con Hipp. | 0.63 | 2.84 | 0.006 |
| 61 | -7 | -2 | cortex right | medial temporal lobe | 256 mm2 | Pt-Con Thal. | 0.55 | -2.48 | 0.015 | Pt-Con Hipp. | 0.62 | 2.79 | 0.007 |
| 65 | -17 | 3 | cortex right | superior temporal gyrus | 244 mm2 | Pt-Con Thal. | 0.51 | -2.27 | 0.026 | Pt-Con Hipp. | 0.77 | 3.48 | 0.001 |
| -36 | -70 | -44 | subcortex | left cerebellum | 200 mm3 | Pt-Con Thal. | 0.54 | 2.43 | 0.017 | Pt-Con Hipp. | 0.77 | -3.48 | 0.001 |

### Characterizing 22q11DS Dysconnectivity Across Thalamic and Hippocampal Sub-Regions

The thalamus and hippocampus are both heterogeneous structures which can be divided into multiple nuclei with distinct physiologies and connectivity profiles (Haber and McFarland, 2001). To assess differential functional connectivity disruptions across thalamic and hippocampal sub-regions, we used a k-means algorithm to cluster thalamic and hippocampal voxels based on unique between-group connectivity differences. The implementation of this algorithm is outlined in **Figure 4**. **Figure 5** shows the k-means solutions for the thalamus and the hippocampus, both of which reveal distinct anterior and posterior clusters. The anterior thalamic cluster encompasses ‘associative’ thalamic nuclei (e.g. the medio-dorsal nucleus), whereas the posterior cluster is centered on visual lateral geniculate and pulvinar nuclei. The hippocampus was similarly divided along an anterior-posterior axis. Seed-based rs-fcMRI was subsequently computed for each thalamic and hippocampal cluster (group contrasts shown in **Figure 5b**). For both the thalamus and hippocampus, the whole-brain connectivity matrices for the anterior cluster were quantitatively more similar to the whole-seed effect (**Figure 5c**, whole-seed effects shown in **Figure 3**). For the thalamus specifically, due to its well-defined neuroanatomical subdivisions in humans, we also investigated how the cluster and whole-seed effects compared to the functional connectivity profiles of seven seeds derived from an FSL diffusion-weighted imaging thalamic atlas (**Figure 6**) (Behrens et al., 2003). As expected, both the anterior thalamic cluster and the whole-thalamus effects were most similar to a set of ‘associative’ thalamic seeds (prefrontal, temporal, premotor). In contrast, the posterior cluster effect was most similar to a set of ‘sensory’ thalamic seeds (occipital, sensory, parietal) (see **Figure 6b**).

**Figure 4.**
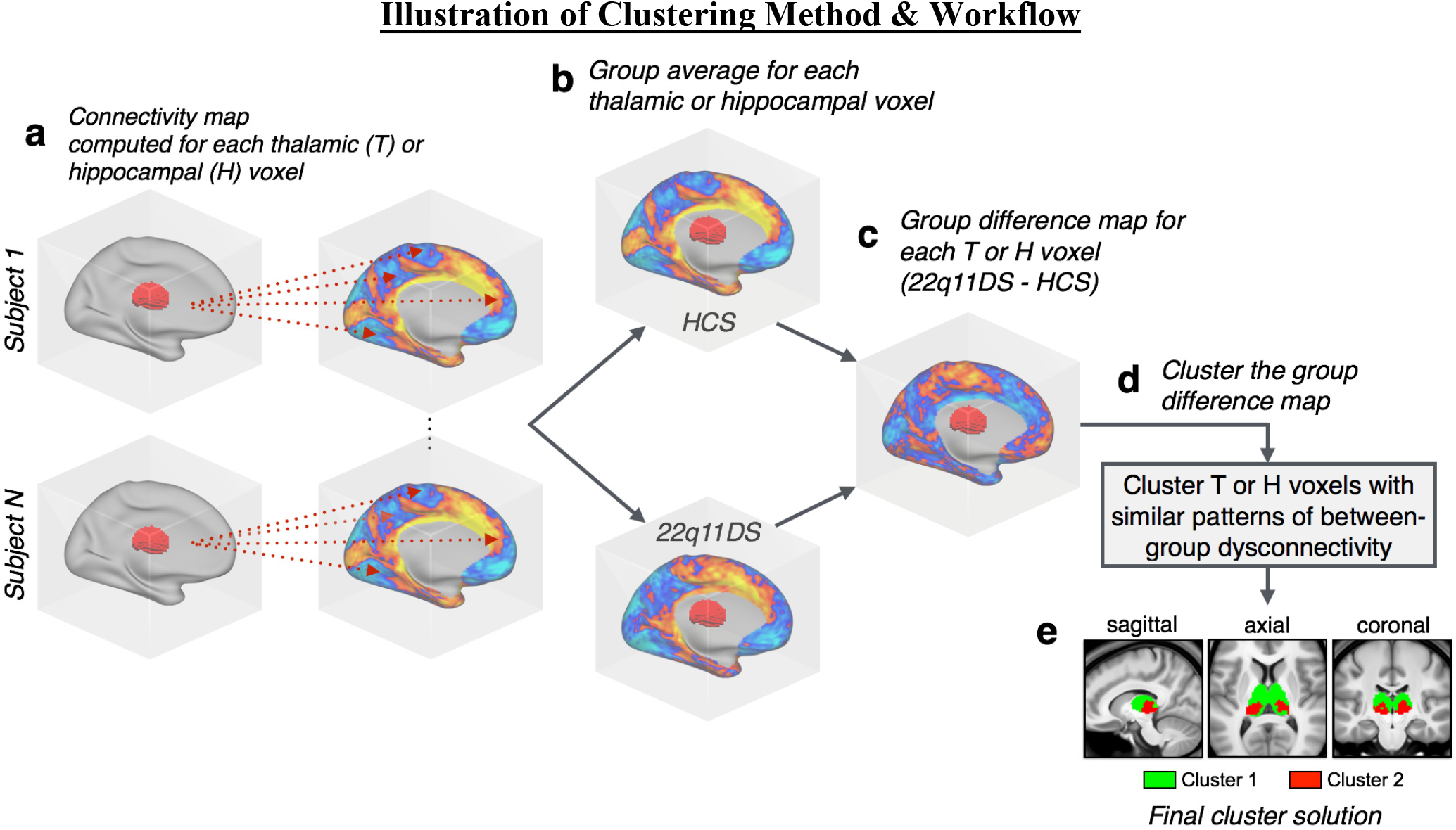
Clustering Algorithm Flow-Chart. A graphical illustration of the procedure for k-means clustering of thalamic voxels. This same procedure was repeated for the hippocampus.

**Figure 5.**
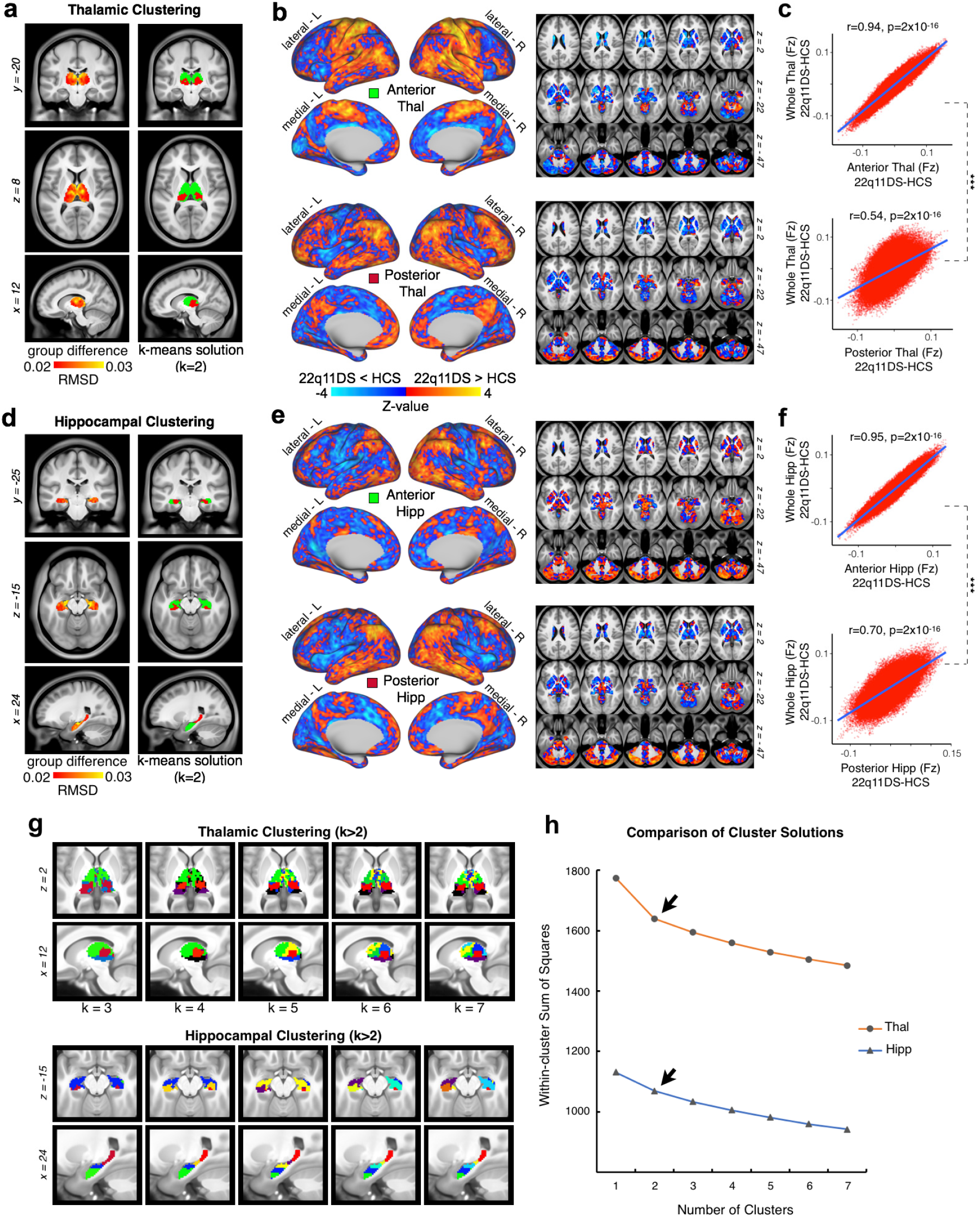
Data-Driven Clustering of Thalamic and Hippocampal Voxels. K-Means clustering of thalamus and hippocampus by group. **(a)** (left) Group difference map (root mean square) for the thalamus, highlighting the anterior and medial dorsal nuclei; (right) the k=2 solution splits the thalamus into an anterior and posterior cluster. **(b)** Surface and volume maps show the group difference (22q11DS v. HCS) for the brain-wide functional connectivity of the anterior thalamic cluster (top), and posterior cluster (bottom). The yellow-orange contrast indicates 22q11DS > HCS. Blue indicates HCS > 22q11DS. (c) Voxel-wise relationship between the brain-wide functional connectivity maps for each thalamic cluster versus the whole-thalamus seed (see **Figure 3**). The correlation with the whole-seed effect is significantly larger for the anterior cluster (top) compared to the posterior cluster (bottom) (Steiger’s z=263, p=2x10^−16^). **(d-f)** Replication of **(a-c)** for the hippocampus, showing a similar anterior/posterior distinction (z=240, p=2x10^−16^). **(g)** Distinct K-means solutions ranging from K=2 to K=7 always reveal an anterior solution (**green**). **(h)** Elbow plot illustrating percent variance explained by each progressive cluster solution.

**Figure 6.**
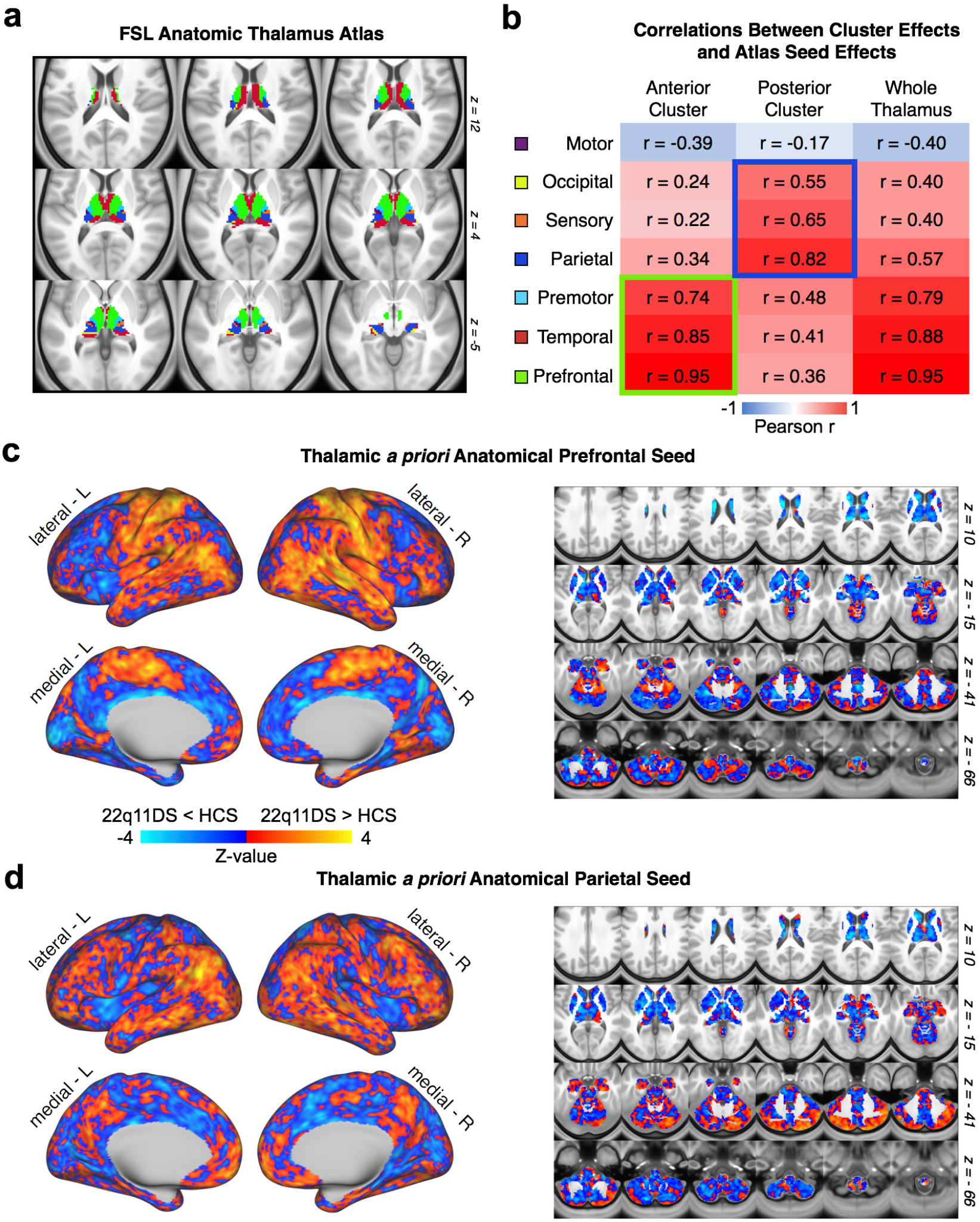
Quantifying Convergence Between K-means Solution and Independent Anatomically-defined Thalamic Seeds. **(a)** FSL anatomic atlas of the thalamus derived from diffusion-weighted imaging. **(b)** Relationship between whole-brain maps of group differences derived from FSL’s atlas and the K=2 K-means solution, indicating a strong correspondence between the ‘anterior’ cluster and executive thalamic nuclei. **(c)** Thalamic *a priori* anatomically-defined prefrontal-projecting thalamic seed used to compute between-group differences, which matches the anterior K=2 effect. **(c)** Thalamic *a priori* anatomically-defined parietal-projecting thalamic seed used to compute between-group differences, which matches the posterior K=2 effect.

### Effects of Global Signal Regression (GSR) on 22q11DS Dysconnectivity Profiles

As noted, prior to the main rs-fcMRI analyses, BOLD data were ‘de-noised’ via mean global signal regression (GSR), in order to attenuate the contribution from spatially pervasive sources of artifact, such as fluctuations in the magnetic field and non-neural physiological processes such as respiration (Power et al., 2017). Nevertheless, there is ongoing development regarding the best-practices for GSR in situations involving clinical populations (Glasser et al., 2017). To test if core observed rs-fcMRI effects are robust to GSR, we re-computed the main analyses without applying GSR. Notably, whole-brain thalamic and hippocampal functional connectivity maps were highly correlated pre- and post-GSR. Furthermore, the pre-GSR data extracted from the original interaction-derived ROIs (see **Figure 2**) showed the same interactive thalamic and hippocampal effects between groups. Finally, the type I error-corrected map for the pre-GSR results (as shown in **Figure 7**) fully overlapped with the original **Figure 2** mask, but with somewhat greater spatial extent (for a detailed list of regions see **Table 5**). As such, results appear robust to GSR.

**Table 5.** Pairwise Comparisons - Region Coordinates, P-values & Effect Sizes (ROIs Indicated Only Pre-GSR)

| X | Y | Z | Hemisphere | Landmark | Size | Comparison | *d* | Mean *T* | *p* | Comparison | *d* | Mean *T* | *p* |
| --- | --- | --- | --- | --- | --- | --- | --- | --- | --- | --- | --- | --- | --- |
| -6 | 15 | 57 | cortex left | superior frontal gyrus | 320 mm2 | Pt-Con Thal. | 0.22 | 0.97 | 0.333 | Pt-Con Hipp. | 0.72 | 3.22 | 0.002 |
| -8 | -35 | -47 | cortex left | marginal sulcus | 536 mm2 | Pt-Con Thal. | 0.01 | 0.04 | 0.966 | Pt-Con Hipp. | 0.59 | 2.66 | 0.010 |
| -8 | -39 | 65 | cortex left | marginal sulcus | 684 mm2 | Pt-Con Thal. | 0.30 | -1.33 | 0.187 | Pt-Con Hipp. | 0.47 | 2.11 | 0.038 |
| -8 | -10 | 47 | cortex left | cingulate sulcus | 652 mm2 | Pt-Con Thal. | 0.01 | 0.06 | 0.954 | Pt-Con Hipp. | 0.68 | 3.04 | 0.003 |
| -7 | -13 | 69 | cortex left | paracentral lobule | 448 mm2 | Pt-Con Thal. | 0.02 | -0.08 | 0.937 | Pt-Con Hipp. | 0.59 | 2.67 | 0.009 |
| -8 | -15 | 65 | cortex left | superior frontal gyrus | 160 mm2 | Pt-Con Thal. | 0.22 | 0.97 | 0.335 | Pt-Con Hipp. | 0.88 | 3.98 | 0.000 |
| -26 | -26 | 70 | cortex left | central sulcus | 584 mm2 | Pt-Con Thal. | 0.03 | -0.15 | 0.880 | Pt-Con Hipp. | 0.74 | 3.34 | 0.001 |
| -35 | -12 | 69 | cortex left | precentral gyrus | 440 mm2 | Pt-Con Thal. | 0.22 | -1.00 | 0.320 | Pt-Con Hipp. | 0.47 | 2.09 | 0.039 |
| -11 | -23 | 75 | cortex left | precentral gyrus | 460 mm2 | Pt-Con Thal. | 0.06 | 0.27 | 0.785 | Pt-Con Hipp. | 0.75 | 3.35 | 0.001 |
| -26 | -1 | 64 | cortex left | superior frontal sulcus | 604 mm2 | Pt-Con Thal. | 0.19 | -0.85 | 0.397 | Pt-Con Hipp. | 0.36 | 1.63 | 0.107 |
| -46 | -46 | 49 | cortex left | intraparietal sulcus | 1408 mm2 | Pt-Con Thal. | 0.32 | -1.46 | 0.149 | Pt-Con Hipp. | 0.36 | 1.62 | 0.109 |
| -35 | -37 | 70 | cortex left | postcentral gyrus | 540 mm2 | Pt-Con Thal. | 0.37 | -1.67 | 0.099 | Pt-Con Hipp. | 0.53 | 2.39 | 0.019 |
| -51 | -25 | 60 | cortex left | postcentral gyrus | 512 mm2 | Pt-Con Thal. | 0.23 | -1.03 | 0.305 | Pt-Con Hipp. | 0.58 | 2.63 | 0.010 |
| -63 | -45 | -4 | cortex left | middle temporal gyrus | 136 mm2 | Pt-Con Thal. | 0.07 | -0.33 | 0.745 | Pt-Con Hipp. | 0.45 | 2.01 | 0.047 |
| -64 | -42 | 15 | cortex left | supramarginal gyrus | 596 mm2 | Pt-Con Thal. | 0.12 | -0.53 | 0.599 | Pt-Con Hipp. | 0.73 | 3.29 | 0.002 |
| -53 | -38 | 16 | cortex left | posterior sylvian fissure | 740 mm2 | Pt-Con Thal. | 0.22 | -1.01 | 0.316 | Pt-Con Hipp. | 0.64 | 2.88 | 0.005 |
| -46 | 4 | 10 | cortex left | insular cortex | 460 mm2 | Pt-Con Thal. | 0.10 | -0.46 | 0.643 | Pt-Con Hipp. | 0.46 | 2.08 | 0.041 |
| -12 | -82 | 39 | cortex left | parieto-occipital sulcus | 620 mm2 | Pt-Con Thal. | 0.27 | -1.20 | 0.233 | Pt-Con Hipp. | 0.62 | 2.81 | 0.006 |
| -7 | -53 | 53 | cortex left | precuneus | 852 mm2 | Pt-Con Thal. | 0.12 | -0.54 | 0.588 | Pt-Con Hipp. | 0.72 | 3.25 | 0.002 |
| -5 | -47 | 34 | cortex left | precuneus | 440 mm2 | Pt-Con Thal. | 0.17 | -0.77 | 0.443 | Pt-Con Hipp. | 0.42 | 1.89 | 0.062 |
| -22 | -95 | 22 | cortex left | cuneus | 408 mm2 | Pt-Con Thal. | 0.07 | -0.31 | 0.759 | Pt-Con Hipp. | 0.51 | 2.30 | 0.024 |
| -35 | -84 | 39 | cortex left | inferior parietal lobule | 84 mm2 | Pt-Con Thal. | 0.06 | -0.27 | 0.791 | Pt-Con Hipp. | 0.44 | 1.98 | 0.051 |
| -27 | -45 | 70 | cortex left | postcentral sulcus | 688 mm2 | Pt-Con Thal. | 0.06 | -0.25 | 0.805 | Pt-Con Hipp. | 0.62 | 2.77 | 0.007 |
| -42 | -91 | 3 | cortex left | inferior occipital gyrus | 192 mm2 | Pt-Con Thal. | 0.31 | -1.39 | 0.168 | Pt-Con Hipp. | 0.27 | 1.23 | 0.221 |
| -46 | -85 | 15 | cortex left | lateral occipital gyrus | 484 mm2 | Pt-Con Thal. | 0.16 | -0.71 | 0.481 | Pt-Con Hipp. | 0.70 | 3.14 | 0.002 |
| -56 | -65 | 8 | cortex left | middle temporal gyrus | 804 mm2 | Pt-Con Thal. | 0.31 | -1.42 | 0.161 | Pt-Con Hipp. | 0.59 | 2.64 | 0.010 |
| -44 | -81 | 38 | cortex left | angular gyrus | 212 mm2 | Pt-Con Thal. | 0.23 | -1.02 | 0.310 | Pt-Con Hipp. | 0.33 | 1.48 | 0.144 |
| -65 | -22 | 28 | cortex left | postcentral sulcus | 864 mm2 | Pt-Con Thal. | 0.16 | -0.71 | 0.480 | Pt-Con Hipp. | 0.58 | 2.59 | 0.011 |
| -60 | -9 | 35 | cortex left | central sulcus | 464 mm2 | Pt-Con Thal. | 0.16 | -0.71 | 0.480 | Pt-Con Hipp. | 0.67 | 3.03 | 0.003 |
| -63 | 0 | 24 | cortex left | central sulcus | 596 mm2 | Pt-Con Thal. | 0.03 | -0.15 | 0.883 | Pt-Con Hipp. | 0.78 | 3.51 | 0.001 |
| -55 | 15 | 3 | cortex left | inferior frontal gyrus | 140 mm2 | Pt-Con Thal. | 0.26 | 1.18 | 0.244 | Pt-Con Hipp. | 0.92 | 4.16 | 0.000 |
| -49 | 39 | 19 | cortex left | middle frontal gyrus | 268 mm2 | Pt-Con Thal. | 0.01 | -0.06 | 0.949 | Pt-Con Hipp. | 0.64 | 2.89 | 0.005 |
| -60 | 8 | 8 | cortex left | inferior frontal gyrus | 208 mm2 | Pt-Con Thal. | 0.29 | 1.30 | 0.196 | Pt-Con Hipp. | 0.94 | 4.22 | 0.000 |
| -56 | 12 | 22 | cortex left | inferior frontal sulcus | 328 mm2 | Pt-Con Thal. | 0.22 | -0.99 | 0.325 | Pt-Con Hipp. | 0.37 | 1.65 | 0.104 |
| -6 | -90 | 13 | cortex left | cuneus | 136 mm2 | Pt-Con Thal. | 0.06 | 0.29 | 0.776 | Pt-Con Hipp. | 0.69 | 3.10 | 0.003 |
| -6 | -84 | 20 | cortex left | cuneus | 304 mm2 | Pt-Con Thal. | 0.08 | -0.35 | 0.730 | Pt-Con Hipp. | 0.63 | 2.82 | 0.006 |
| -12 | -75 | 14 | cortex left | cuneus | 380 mm2 | Pt-Con Thal. | 0.05 | -0.24 | 0.808 | Pt-Con Hipp. | 0.75 | 3.36 | 0.001 |
| -15 | -63 | 2 | cortex left | calcarine sulcus | 612 mm2 | Pt-Con Thal. | 0.06 | -0.26 | 0.796 | Pt-Con Hipp. | 0.86 | 3.85 | 0.000 |
| -46 | 12 | 52 | cortex left | middle frontal gyrus | 624 mm2 | Pt-Con Thal. | 0.08 | -0.35 | 0.729 | Pt-Con Hipp. | 0.62 | 2.77 | 0.007 |
| -46 | -14 | -1 | cortex left | medial temporal lobe | 448 mm2 | Pt-Con Thal. | 0.10 | -0.44 | 0.664 | Pt-Con Hipp. | 0.60 | 2.71 | 0.008 |
| -51 | 6 | -18 | cortex left | medial temporal lobe | 256 mm2 | Pt-Con Thal. | 0.53 | -2.38 | 0.020 | Pt-Con Hipp. | 0.36 | 1.63 | 0.107 |
| -55 | -19 | 3 | cortex left | medial temporal lobe | 476 mm2 | Pt-Con Thal. | 0.12 | 0.56 | 0.576 | Pt-Con Hipp. | 1.00 | 4.50 | 0.000 |
| -58 | 3 | -14 | cortex left | superior temporal gyrus | 224 mm2 | Pt-Con Thal. | 0.01 | 0.03 | 0.978 | Pt-Con Hipp. | 0.75 | 3.38 | 0.001 |
| -63 | -15 | 0 | cortex left | superior temporal gyrus | 352 mm2 | Pt-Con Thal. | 0.02 | -0.07 | 0.942 | Pt-Con Hipp. | 0.90 | 4.03 | 0.000 |
| -61 | -11 | -20 | cortex left | superior temporal sulcus | 244 mm2 | Pt-Con Thal. | 0.18 | -0.81 | 0.420 | Pt-Con Hipp. | 0.50 | 2.23 | 0.029 |
| -61 | -27 | -11 | cortex left | superior temporal sulcus | 220 mm2 | Pt-Con Thal. | 0.01 | 0.04 | 0.968 | Pt-Con Hipp. | 0.70 | 3.14 | 0.002 |
| 13 | -54 | 3 | cortex right | isthmus | 580 mm2 | Pt-Con Thal. | 0.06 | -0.25 | 0.802 | Pt-Con Hipp. | 0.64 | 2.88 | 0.005 |
| 15 | -58 | -5 | cortex right | parieto-occipital sulcus | 376 mm2 | Pt-Con Thal. | 0.21 | -0.95 | 0.344 | Pt-Con Hipp. | 0.51 | 2.30 | 0.024 |
| 7 | 2 | 48 | cortex right | cingulate sulcus | 504 mm2 | Pt-Con Thal. | 0.01 | 0.05 | 0.962 | Pt-Con Hipp. | 0.61 | 2.74 | 0.008 |
| 6 | 16 | 39 | cortex right | cingulate sulcus | 248 mm2 | Pt-Con Thal. | 0.24 | 1.06 | 0.291 | Pt-Con Hipp. | 0.71 | 3.19 | 0.002 |
| 3 | 6 | 36 | cortex right | cingulate gyrus | 132 mm2 | Pt-Con Thal. | 0.35 | 1.57 | 0.120 | Pt-Con Hipp. | 1.05 | 4.72 | 0.000 |
| 8 | -48 | 53 | cortex right | precuneus | 664 mm2 | Pt-Con Thal. | 0.08 | -0.36 | 0.718 | Pt-Con Hipp. | 0.62 | 2.80 | 0.006 |
| 11 | -36 | 75 | cortex right | postcentral gyrus | 604 mm2 | Pt-Con Thal. | 0.00 | -0.01 | 0.996 | Pt-Con Hipp. | 0.75 | 3.36 | 0.001 |
| 6 | -23 | 59 | cortex right | paracentral lobule | 424 mm2 | Pt-Con Thal. | 0.28 | -1.27 | 0.209 | Pt-Con Hipp. | 0.33 | 1.47 | 0.147 |
| 6 | -8 | 61 | cortex right | paracentral lobule | 380 mm2 | Pt-Con Thal. | 0.07 | -0.31 | 0.756 | Pt-Con Hipp. | 0.50 | 2.25 | 0.027 |
| 40 | -18 | 62 | cortex right | precentral gyrus | 312 mm2 | Pt-Con Thal. | 0.31 | -1.38 | 0.171 | Pt-Con Hipp. | 0.52 | 2.35 | 0.021 |
| 44 | -3 | 61 | cortex right | precentral gyrus | 120 mm2 | Pt-Con Thal. | 0.09 | 0.42 | 0.675 | Pt-Con Hipp. | 0.53 | 2.39 | 0.019 |
| 28 | -17 | 74 | cortex right | precentral gyrus | 408 mm2 | Pt-Con Thal. | 0.18 | -0.79 | 0.430 | Pt-Con Hipp. | 0.68 | 3.07 | 0.003 |
| 12 | -14 | 71 | cortex right | precentral sulcus | 396 mm2 | Pt-Con Thal. | 0.10 | -0.44 | 0.663 | Pt-Con Hipp. | 0.50 | 2.24 | 0.028 |
| 23 | 15 | 65 | cortex right | superior frontal sulcus | 356 mm2 | Pt-Con Thal. | 0.46 | -2.08 | 0.041 | Pt-Con Hipp. | 0.13 | 0.57 | 0.572 |
| 32 | -30 | 71 | cortex right | postcentral gyrus | 488 mm2 | Pt-Con Thal. | 0.10 | -0.43 | 0.670 | Pt-Con Hipp. | 0.82 | 3.67 | 0.000 |
| 42 | -34 | 0 | cortex right | central sulcus | 460 mm2 | Pt-Con Thal. | 0.36 | -1.63 | 0.107 | Pt-Con Hipp. | 0.52 | 2.35 | 0.021 |
| 50 | -14 | 54 | cortex right | postcentral gyrus | 248 mm2 | Pt-Con Thal. | 0.28 | -1.25 | 0.216 | Pt-Con Hipp. | 0.43 | 1.93 | 0.058 |
| 64 | -38 | 0 | cortex right | middle temporal gyrus | 280 mm2 | Pt-Con Thal. | 0.24 | -1.08 | 0.285 | Pt-Con Hipp. | 0.41 | 1.84 | 0.069 |
| 62 | -46 | 24 | cortex right | superior temporal sulcus | 664 mm2 | Pt-Con Thal. | 0.02 | -0.10 | 0.921 | Pt-Con Hipp. | 0.66 | 2.96 | 0.004 |
| 68 | -30 | 9 | cortex right | superior temporal gyrus | 376 mm2 | Pt-Con Thal. | 0.16 | -0.73 | 0.469 | Pt-Con Hipp. | 0.58 | 2.61 | 0.011 |
| 57 | -23 | 10 | cortex right | medial temporal lobe | 432 mm2 | Pt-Con Thal. | 0.02 | 0.09 | 0.930 | Pt-Con Hipp. | 0.90 | 4.05 | 0.000 |
| 56 | -32 | -1 | cortex right | medial temporal lobe | 540 mm2 | Pt-Con Thal. | 0.02 | 0.09 | 0.930 | Pt-Con Hipp. | 0.72 | 3.22 | 0.002 |
| 44 | -23 | 16 | cortex right | insular cortex | 572 mm2 | Pt-Con Thal. | 0.07 | 0.30 | 0.765 | Pt-Con Hipp. | 0.71 | 3.21 | 0.002 |
| 49 | 48 | 5 | cortex right | inferior frontal gyrus | 556 mm2 | Pt-Con Thal. | 0.22 | -1.01 | 0.318 | Pt-Con Hipp. | 0.60 | 2.71 | 0.008 |
| 45 | -5 | -2 | cortex right | insular cortex | 316 mm2 | Pt-Con Thal. | 0.09 | -0.39 | 0.696 | Pt-Con Hipp. | 0.68 | 3.08 | 0.003 |
| 44 | 10 | -8 | cortex right | insular cortex | 68 mm2 | Pt-Con Thal. | 0.13 | 0.59 | 0.558 | Pt-Con Hipp. | 0.61 | 2.75 | 0.007 |
| 45 | 29 | -1 | cortex right | orbital gyrus | 604 mm2 | Pt-Con Thal. | 0.14 | -0.65 | 0.517 | Pt-Con Hipp. | 0.78 | 3.51 | 0.001 |
| 8 | -67 | 28 | cortex right | parieto-occipital sulcus | 276 mm2 | Pt-Con Thal. | 0.17 | -0.78 | 0.440 | Pt-Con Hipp. | 0.52 | 2.32 | 0.023 |
| 18 | -59 | 63 | cortex right | precuneus | 652 mm2 | Pt-Con Thal. | 0.24 | -1.08 | 0.284 | Pt-Con Hipp. | 0.34 | 1.53 | 0.131 |
| 21 | -76 | 47 | cortex right | parieto-occipital sulcus | 476 mm2 | Pt-Con Thal. | 0.23 | -1.05 | 0.298 | Pt-Con Hipp. | 0.44 | 1.96 | 0.053 |
| 34 | -51 | 43 | cortex right | intraparietal sulcus | 724 mm2 | Pt-Con Thal. | 0.28 | -1.24 | 0.217 | Pt-Con Hipp. | 0.24 | 1.08 | 0.284 |
| 29 | -70 | 53 | cortex right | superior parietal lobule | 144 mm2 | Pt-Con Thal. | 0.16 | -0.71 | 0.477 | Pt-Con Hipp. | 0.29 | 1.29 | 0.201 |
| 47 | -59 | 19 | cortex right | angular gyrus | 820 mm2 | Pt-Con Thal. | 0.28 | -1.26 | 0.212 | Pt-Con Hipp. | 0.44 | 1.98 | 0.051 |
| 62 | -34 | 47 | cortex right | supramarginal gyrus | 80 mm2 | Pt-Con Thal. | 0.34 | -1.53 | 0.129 | Pt-Con Hipp. | 0.05 | 0.21 | 0.835 |
| 65 | -14 | 18 | cortex right | postcentral gyrus | 700 mm2 | Pt-Con Thal. | 0.06 | -0.26 | 0.792 | Pt-Con Hipp. | 0.64 | 2.87 | 0.005 |
| 53 | -25 | 40 | cortex right | postcentral sulcus | 820 mm2 | Pt-Con Thal. | 0.40 | -1.78 | 0.079 | Pt-Con Hipp. | 0.47 | 2.13 | 0.036 |
| 65 | -1 | 25 | cortex right | central sulcus | 684 mm2 | Pt-Con Thal. | 0.15 | -0.68 | 0.498 | Pt-Con Hipp. | 0.72 | 3.22 | 0.002 |
| 59 | -11 | 44 | cortex right | postcentral gyrus | 508 mm2 | Pt-Con Thal. | 0.28 | -1.28 | 0.205 | Pt-Con Hipp. | 0.58 | 2.60 | 0.011 |
| 52 | 34 | 22 | cortex right | middle frontal gyrus | 620 mm2 | Pt-Con Thal. | 0.28 | -1.25 | 0.215 | Pt-Con Hipp. | 0.41 | 1.86 | 0.067 |
| 60 | 10 | 7 | cortex right | precentral gyrus | 304 mm2 | Pt-Con Thal. | 0.14 | 0.64 | 0.526 | Pt-Con Hipp. | 0.76 | 3.43 | 0.001 |
| 55 | 21 | 20 | cortex right | inferior frontal sulcus | 572 mm2 | Pt-Con Thal. | 0.29 | -1.31 | 0.193 | Pt-Con Hipp. | 0.56 | 2.53 | 0.013 |
| 20 | 31 | -19 | cortex right | orbital gyrus | 104 mm2 | Pt-Con Thal. | 0.10 | -0.46 | 0.644 | Pt-Con Hipp. | 0.45 | 2.04 | 0.045 |
| 6 | -92 | 9 | cortex right | cuneus | 44 mm2 | Pt-Con Thal. | 0.09 | -0.42 | 0.676 | Pt-Con Hipp. | 0.34 | 1.52 | 0.132 |
| 11 | -78 | 4 | cortex right | calcarine sulcus | 176 mm2 | Pt-Con Thal. | 0.04 | 0.18 | 0.860 | Pt-Con Hipp. | 0.60 | 2.72 | 0.008 |
| 6 | -80 | 31 | cortex right | cuneus | 384 mm2 | Pt-Con Thal. | 0.03 | -0.12 | 0.904 | Pt-Con Hipp. | 0.75 | 3.36 | 0.001 |
| 23 | -63 | 16 | cortex right | parieto-occipital sulcus | 448 mm2 | Pt-Con Thal. | 0.00 | 0.02 | 0.985 | Pt-Con Hipp. | 0.63 | 2.85 | 0.006 |
| 28 | 64 | 12 | cortex right | middle frontal gyrus | 368 mm2 | Pt-Con Thal. | 0.04 | 0.16 | 0.873 | Pt-Con Hipp. | 0.63 | 2.84 | 0.006 |
| 7 | 66 | 1 | cortex right | superior frontal gyrus | 204 mm2 | Pt-Con Thal. | 0.21 | -0.94 | 0.350 | Pt-Con Hipp. | 0.42 | 1.88 | 0.064 |
| 8 | 48 | 6 | cortex right | anterior cingulate cortex | 352 mm2 | Pt-Con Thal. | 0.02 | 0.08 | 0.935 | Pt-Con Hipp. | 0.71 | 3.17 | 0.002 |
| 15 | 65 | 22 | cortex right | superior frontal gyrus | 248 mm2 | Pt-Con Thal. | 0.11 | -0.51 | 0.614 | Pt-Con Hipp. | 0.52 | 2.33 | 0.022 |
| 30 | 50 | 36 | cortex right | superior frontal gyrus | 508 mm2 | Pt-Con Thal. | 0.22 | -0.97 | 0.334 | Pt-Con Hipp. | 0.39 | 1.77 | 0.081 |
| 54 | 5 | 49 | cortex right | precentral gyrus | 460 mm2 | Pt-Con Thal. | 0.25 | -1.13 | 0.261 | Pt-Con Hipp. | 0.40 | 1.78 | 0.078 |
| 46 | 8 | -17 | cortex right | medial temporal lobe | 224 mm2 | Pt-Con Thal. | 0.36 | -1.64 | 0.105 | Pt-Con Hipp. | 0.52 | 2.33 | 0.022 |
| 54 | 8 | -16 | cortex right | superior temporal gyrus | 112 mm2 | Pt-Con Thal. | 0.35 | -1.56 | 0.122 | Pt-Con Hipp. | 0.39 | 1.77 | 0.081 |
| 67 | -13 | 5 | cortex right | superior temporal gyrus | 268 mm2 | Pt-Con Thal. | 0.10 | -0.46 | 0.647 | Pt-Con Hipp. | 0.71 | 3.18 | 0.002 |
| 59 | 0 | -20 | cortex right | superior temporal sulcus | 336 mm2 | Pt-Con Thal. | 0.38 | -1.72 | 0.090 | Pt-Con Hipp. | 0.37 | 1.67 | 0.099 |

**Figure 7.**
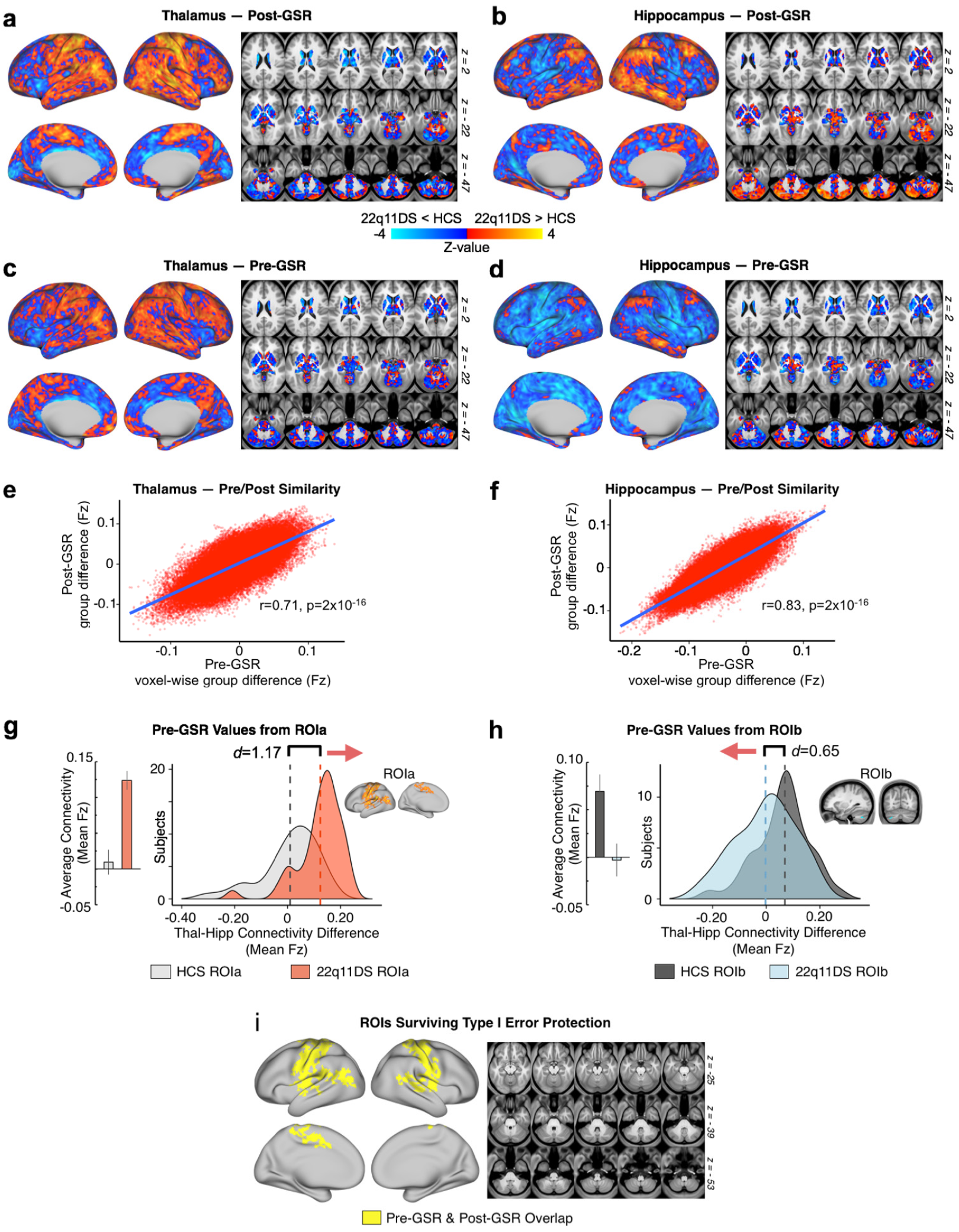
Stability of Effects Before and After Global Signal Regression. Here we show a comparison of pre- and post-GSR effects. **(a-b)** Post-GSR threshold-free connectivity for thalamus and hippocampus (same as **Figure 3**). **(c-d)** Thalamic and hippocampal connectivity before GSR. **(e-f)** Pearson correlation between pre- and post-GSR matrices. **(g-h)** pre-GSR data extracted from ROIa and ROIb (see **Figure 2**). **(i)** Overlapping regions (logical AND) for type I error-corrected interaction effect pre-GSR and post-GSR. The cerebellar effect (**Figure 2** ROIb) did not survive without GSR.

### Interactive 22q11DS Disruptions across Sensory and Executive Networks

As noted, while we observed a focused type I error corrected effect in the cerebellum, the interactive results appeared substantially more widespread. Therefore, we tested whether 22q11DS patients indeed exhibit a network-level dissociation for thalamic versus hippocampal connectivity. To this end, we repeated the seed-based analyses focusing on thalamic and hippocampal connectivity to *a priori* networks derived from a data-driven parcellation of the human cortex, cerebellum, and striatum (Buckner et al., 2011; Choi et al., 2012; Yeo et al., 2011) (**Figure 8**). Here, functional connectivity was computed between the thalamic and hippocampal seeds and each of the seven *a priori* networks. In other words, we examined the connectivity between the thalamus or hippocampus with the entire brain-wide average of each functional network, yielding 14 values (i.e. 7 thalamus-to-each-network and 7 hippocampus-to-each-network rs-fcMRI values). As predicted, the 22q11DS group exhibited significantly increased thalamic but decreased hippocampal connectivity to brain-wide somatomotor (SOM) network regions, while the opposite effect was observed for the brain-wide frontoparietal (FPN) network regions.

**Figure 8.**
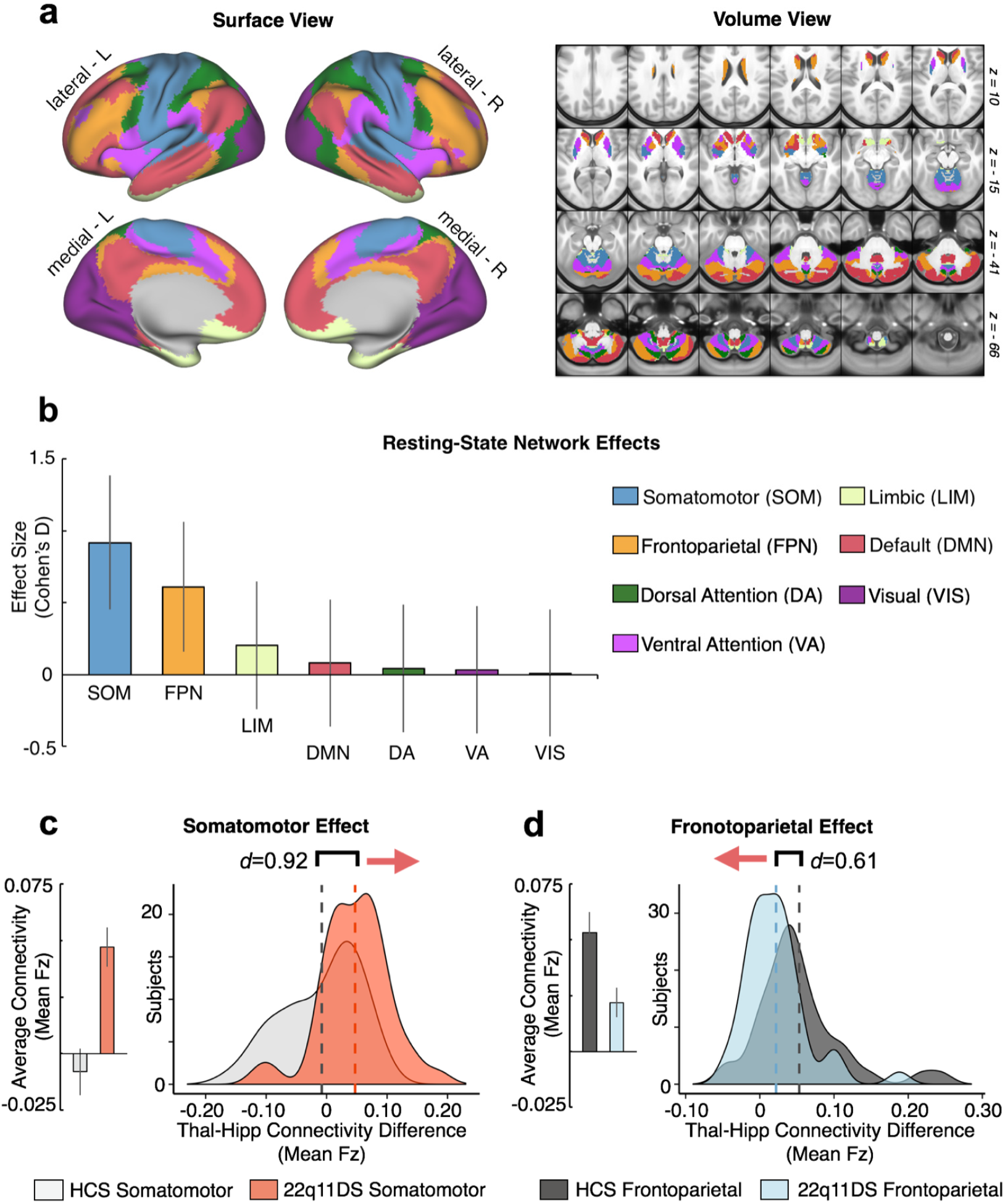
Examining Effects Across *a priori* Functional Networks. Replication of seed-based analysis (**Figure 2**) using *a priori* functionally-derived networks, mapped into CIFTI space **(a)**. Surface and volume components of the map showing seven distinct functional networks derived from prior work that parcellated the cortex, striatum and cerebellum. Colors indicate distinct functional networks, following the same labeling pattern as the original work. **(b)** Effect sizes (Cohen’s D) with 95% confidence intervals comparing 22q11DS and HCS groups with regard to the difference scores between thalamic and hippocampal connectivity to each of the seven networks. **(c)** Thalamus-hippocampus difference scores illustrated for the somatomotor network (SOM) across subjects in each group. Group means (left) and distributions (right) illustrate the direction of the effect. **(d)** Same as **(c)** for the frontoparietal network (FPN), showing an effect in the opposite direction as SOM.

Critically, across subjects, for both the interaction-derived and *a priori* network-derived effects, the magnitude of the rs-fcMRI effect in the sensory ROI/network was inversely related to the magnitude of the effect in the associative ROI/network (**Figures 9** and **10**). We quantified this relationship via Pearson correlations between the effects defined in the data-driven interaction-derived ROIs and the *a priori* networks (SOM and FPN).

**Figure 9.**
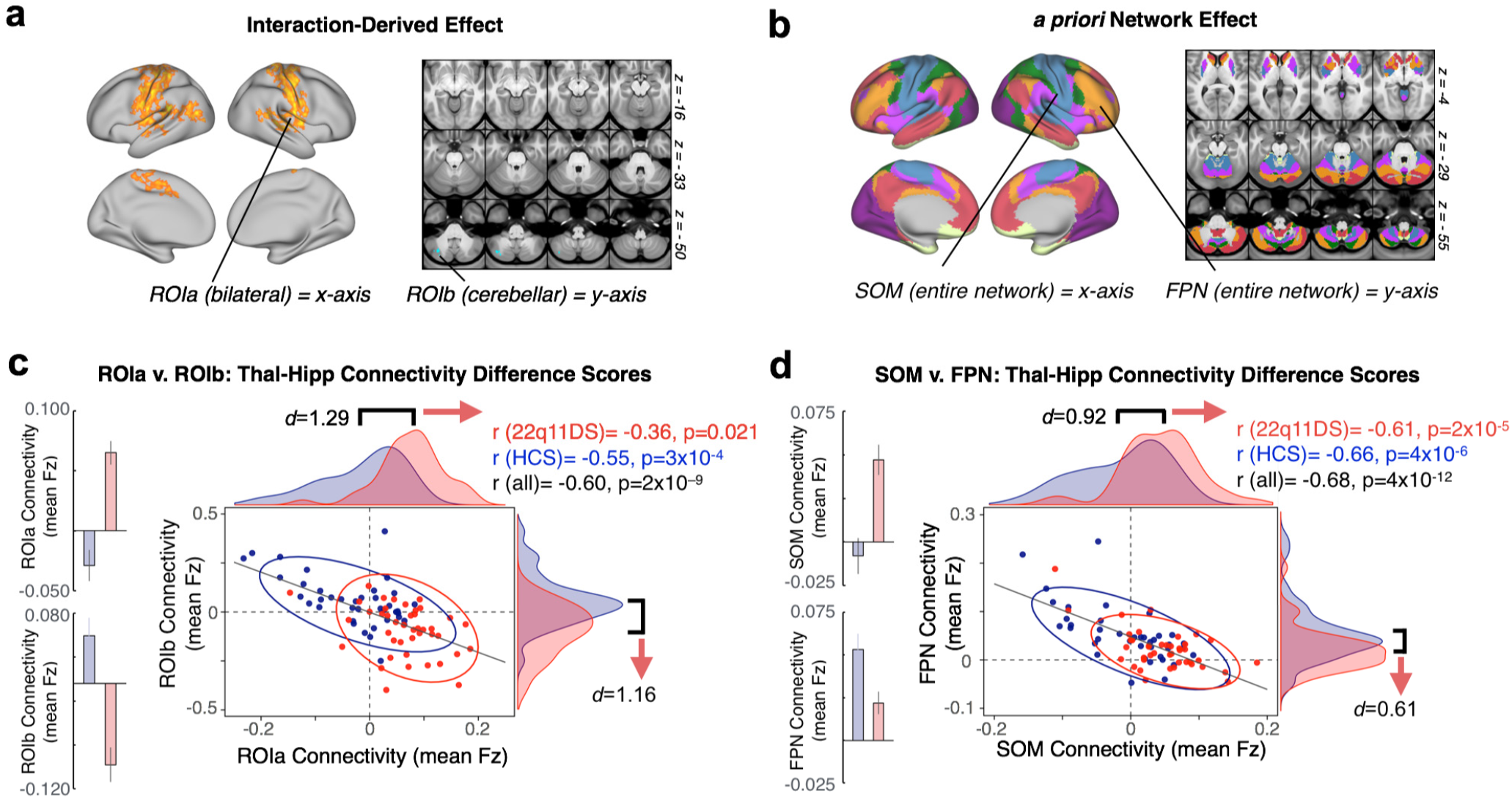
Relationship Between Regions of Reciprocally Disrupted Connectivity. **(a)** Across subjects, ROIa and ROIb (see **Figure 2**) are significantly negatively related in terms of the overall connectivity effect (thalamus-hippocampus Fz difference score). The distribution of 22q11DS subjects (red) is distinctly shifted relative to controls (blue), showing greater thalamic connectivity relative to hippocampal connectivity for ROIa, and the inverse for ROIb. **(b)** Replication of **(a)** using connectivity to *a priori* somatomotor and frontoparietal networks (see **Figure 8**), demonstrating that these reciprocal effects map onto large-scale sensory and associative networks. 22q11DS subjects show increased thalamic and decreased hippocampal connectivity to sensory regions, but the opposite effect in associative regions. Note: effects are presented for each seed and ROI pair individually in **Figure 10**.

**Figure 10.**
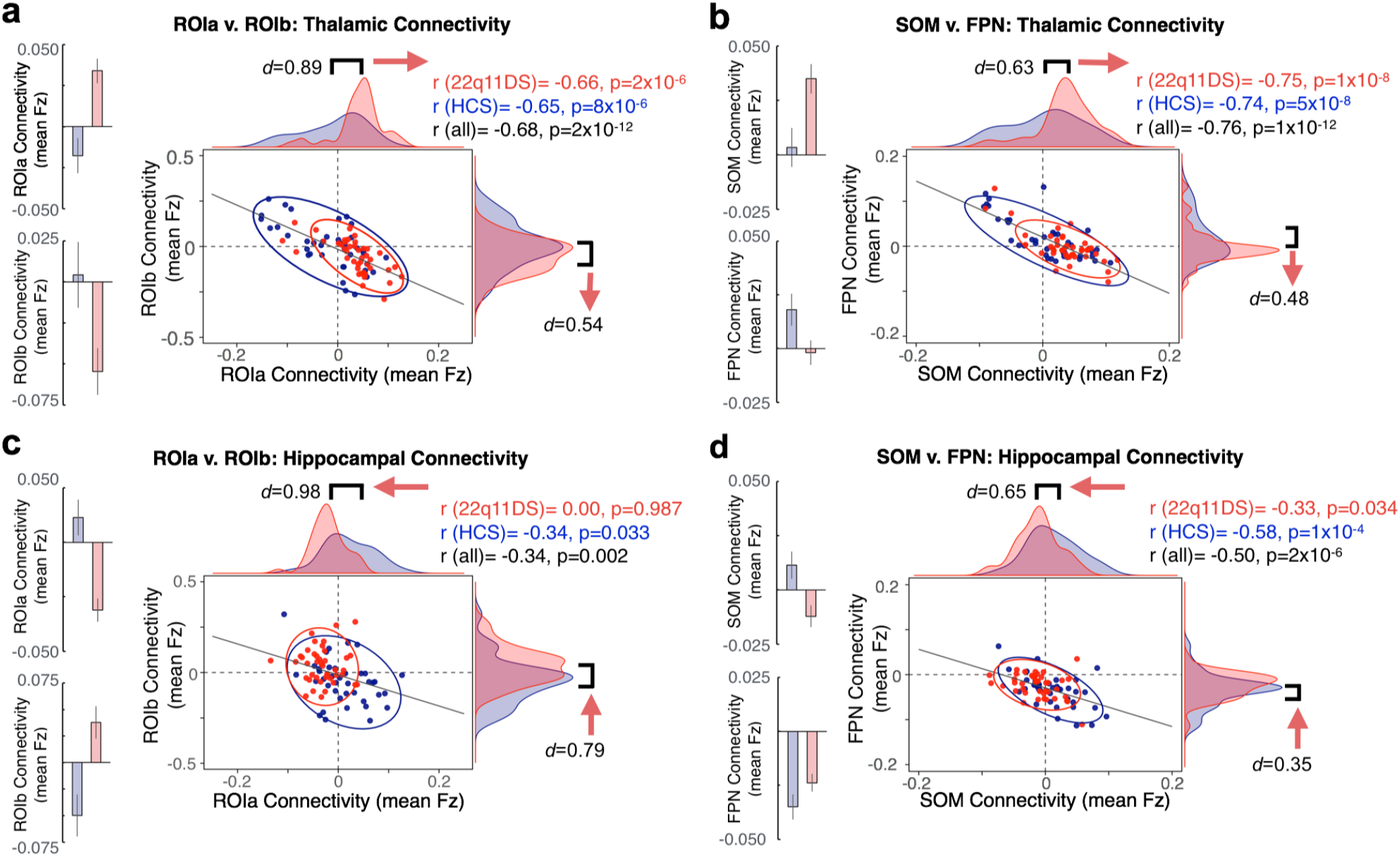
Reciprocal Effects Across Thalamic and Hippocampal Seeds. Expanding on **Figure 9**, showing distributions and relationships across subjects for whole thalamic and hippocampal seeds individually. The thalamic effect shifts in the opposite direction to the hippocampal effect.

### Prediction of 22q11 Case-Control Status from Data-driven and Network-level Dysconnectivity Effects

To test the hypothesis that the observed rs-fcMRI effects have potential utility as a neural biomarker, we conducted a SVM analysis (**Figure 11**). One-dimensional SVMs, computed based on the unweighted linear combination of thalamic and hippocampal connectivity to ROIa and ROIb (interaction-derived ROIs) correctly predicted diagnosis at rates well above chance (for n=1000 iterations, mean AUC=0.843, SD=0.043). The unweighted combination of thalamic and hippocampal connectivity to entire *a priori* SOM and FPN networks was also able to provide moderate diagnostic accuracy (for n=1000 iterations, mean AUC=0.739, SD=0.057). The four-dimensional SVM solution (i.e. combing the four discovered features: thal-ROIa, thal-ROIb, hipp-ROIa, hipp-ROIb), which separated the groups by attempting to fit a hyper-plane that optimally weighted each factor (thalamic and hippocampal connectivity to each ROI/network), provided no performance advantage relative to the unweighted combination of features.

**Figure 11.**
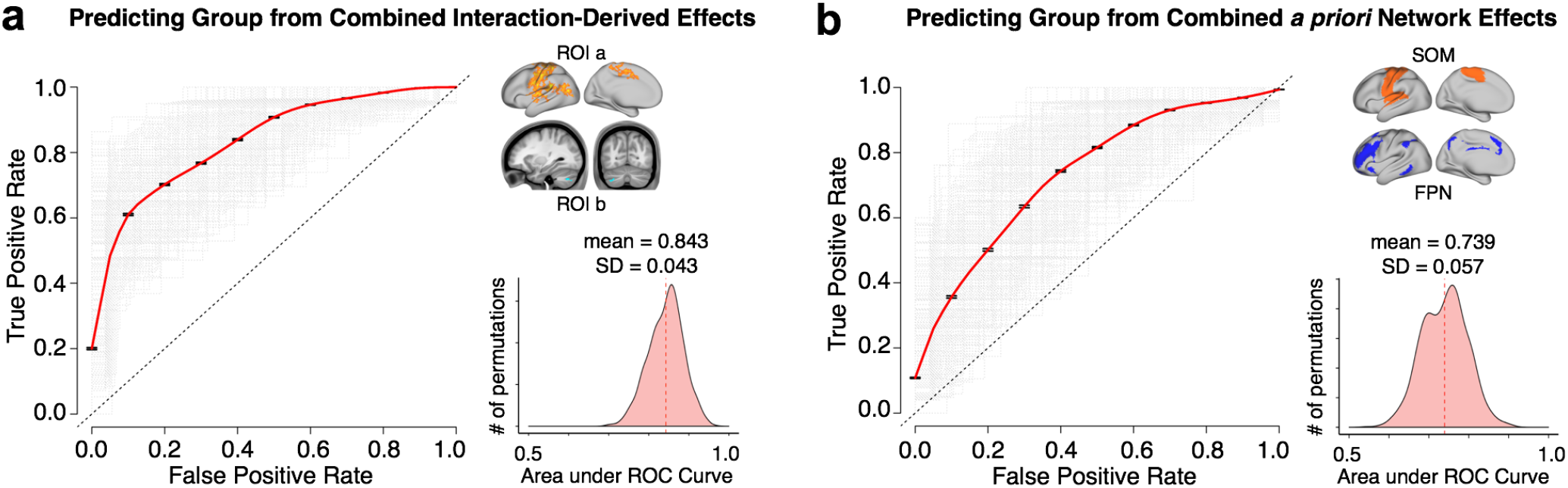
Diagnostic Classification via Support Vector Machine. SVM classification based on combined fc-MRI effects. **(a)** ROC curves for a binary classifier predicting group membership (22q11DS or HCS) based on the linear combination of connectivity values (mean Fz) from the thalamus and hippocampus to ROIa and ROIb ([thalROIa + hippROIb] - [thalROIb + hippROIa]). ROC curves for each of the n=1000 iterations are plotted in gray, and the vertical average in red. The distribution of areas under the n=1000 ROC curves (AUC) is plotted on the right, where an AUC of 1 would represent a perfect classifier. **(b)** replication of **(a)** using the linear combination of thalamic and hippocampal connectivity to somatomotor and frontoparietal networks ([thalSOM + hippFPN] - [thalFPN + hippSOM]).

## DISCUSSION

22q11DS is associated with notable neural alterations and presents a compelling genetic high-risk model in which anomalous circuitry can be investigated prior to development of overt psychiatric illness. Yet, there is a knowledge gap in our understanding of translational neural phenotypes in a genetic risk model such as 22q11DS. The thalamo-cortical system presents a unique leverage point for investigations of brain-wide dysconnectivity given its central locus and key role in functional and structural loops across the brain (Behrens et al., 2003; Zhang et al., 2010). Similarly, the hippocampus exhibits distinct brain-wide rs-fcMRI patterns relative to the thalamus in healthy humans (**Figure 1**), and structural and functional hippocampal alterations feature prominently across the neuropsychiatric spectrum (Tamminga et al., 2010). Notably, disruptions of this circuitry have been identified in a mouse model of the 22q11.2 deletion (Sigurdsson et al., 2010; Chun et al., 2014). The present study, for the first time, identified opposing patterns of thalamic-hippocampal disruption in human 22q11.2 deletion carriers. Specifically, findings revealed a pattern of significant thalamic overconnectivity with bilateral sensorimotor regions, including auditory cortex, in 22q11DS relative to typically developing controls, with the opposite effect in cerebellar regions. These findings extend prior work by identifying reciprocal and functionally linked disruptions of hippocampal connectivity in 22q11DS. This effect was verified via *a priori* networks derived from a data-driven parcellation of the human cortex, cerebellum, and striatum (Buckner et al., 2011; Yeo et al., 2011; Choi et al., 2012). Again, the 22q11DS group showed an inverse pattern relative to controls, with significantly increased thalamic and decreased hippocampal connectivity to SOM regions, and the opposite effect in FPN networks. Data-driven k-means clustering showed, for the first time, that the anterior portions of thalamus and hippocampus were driving the observed patterns of disrupted connectivity. This result is in concert with the view that these are heterogeneous structures, which can be divided into multiple nuclei with distinct physiologies and connectivity profiles. Finally, machine learning analyses revealed accurate classification of 22q11DS patients versus controls, based on the un-weighted linear combination of thalamic and hippocampal connectivity at rates well above chance (84% overall classification accuracy). These effects indicate potential utility of the reported thalamo-hippocampal dysconnectivity for prediction of future development of neuropsychiatric symptoms.

### Implications for the Neurobiology of Psychosis

Thalamic over-connectivity with sensorimotor regions and cerebellar under-connectivity in 22q11DS are in line with prior observations in patients with established schizophrenia (Anticevic et al., 2014) and those at clinical high-risk for the disorder (Anticevic et al., 2015b). Notably, the reported thalamic effect was particularly prominent in those clinical high-risk youth who subsequently converted to psychosis, which would suggest that these network-level disturbances are present prior to onset of overt illness. Note that in 22q11DS, hypo-connectivity with broader executive regions was supported by network-level FPN analysis. Our findings of reciprocal thalamic-hippocampal effects are particularly notable, given that the nucleus reuniens of the thalamus directly innervates the hippocampus (Herkenham, 1978; Lisman, 2012), and was recently determined to play a key role in regulating bi-directional communication between the dorsal hippocampus and medial prefrontal cortex (Hallock et al., 2016). This hypothesis is further supported by the k-means solutions, which implicate the key functional roles of ‘anterior’ subdivisions for both thalamic and hippocampal seeds.

Nevertheless, BOLD rs-fMRI is an indirect observational neuroimaging measure, and thus cannot address underlying cellular mechanisms. However, these processes can be investigated in translational studies in animal (Hiroi et al., 2013), and *in vitro* models (Brennand et al., 2012) as well as computational modeling studies, which can generate testable predictions at the circuit level (Anticevic et al., 2015a). Theoretical models of psychosis implicate alterations in glutamatergic, dopaminergic and inhibitory GABAergic neurotransmission, which may be relevant to the observed disruptions of thalamo-striatal-cortical circuitry (Gonzalez-Burgos and Lewis, 2012; Lewis et al., 2012; Woodward et al., 2012). At present, the origin of the widespread reciprocal thalamic-hippocampal disruption is not fully understood. However, investigation of this circuitry in the context of a well-characterized genetic etiology, as demonstrated in the current study, is a key advantage and a path forward. One possibility may involve dysfunction of N-methyl-D-aspartate glutamate receptors (NMDAR) (Javitt, 2007; Loh et al., 2007), which may impact excitatory-inhibitory balance in cortical circuits and lead to large-scale disturbances in thalamo-cortical information flow. Notably, this hypothesis is supported by data from the 22q11.2 mouse model, as discussed below. Alternatively, it is possible that a local ‘hotspot’ of dysfunction (e.g. such as the nucleus reuniens of the thalamus) emerges, via confluence of polygenic risk (Anticevic and Lisman, 2017).

### Convergence with 22q11.2 Mouse Model

In a mouse model of the 22q11.2 deletion, Chun and colleagues reported disrupted glutamatergic synaptic transmission at thalamic inputs to the auditory cortex (Chun et al., 2014), suggesting that thalamo-cortical disruption could be a pathogenic mechanism that mediates susceptibility to positive psychotic symptoms in 22q11DS. Furthermore, it was determined that thalamo-cortical disruption in 22q11DS mice was caused by abnormal elevation of dopamine D2 (DRD2) receptors in the thalamus. Increased DRD2 in the thalamus and other brain regions has been reported in antipsychotic naïve schizophrenia patients (Oke et al., 1988; Cronenwett and Csernansky, 2010). The *dgcr8* gene, which encodes part of the microprocessor complex that mediates microRNA (miRNA) biogenesis, was pinpointed as being responsible for this neuronal phenotype in the 22q11DS mouse model. Consequently, reduced dosage of *dgcr8* in 22q11DS may lead to miRNA dysregulation, and downstream disruption of synaptic function and proper neural circuit development (Earls and Zakharenko, 2014).

More recently, Chun et al. further established a thalamus-enriched miRNA in the 22q11DS mouse model, which specifically targets DRD2 (miRNA 338-3p). This may be a key mediator of the disruption of synaptic transmission at thalamo-cortical projections and the late adolescent/early adult onset of auditory perceptual anomalies in individuals with 22q11DS (Chun et al., 2016).

Although, to our knowledge, reciprocal disruption of the hippocampal-thalamic circuit has not yet been directly probed in this mouse model, there is complementary evidence for impaired synchronization of neural activity between the hippocampus and prefrontal cortex. Specifically, Sigurdsson and colleagues (2010) found that, while hippocampal-prefrontal synchrony increased during working memory performance in wild-type mice, this phase-locking did not occur in the 22q11DS mice. Further, the magnitude of baseline hippocampal-prefrontal coherence was predictive of how long it took the mice to learn the task. Taken together, these findings suggest that observations of disrupted large-scale network coherence in human 22q11.2 deletion carriers are recapitulated in the animal model. Recent studies from rodent models have also revealed a broader role of the thalamus in higher-order cognitive functions such as working memory. In fact, working memory maintenance *required* mediodorsal thalamic inputs, suggesting a causal role of dysfunction in this circuit in characteristic cognitive deficits associated with schizophrenia (Bolkan et al., 2017).

### Pitfalls and Future Solutions

Notably, only a minority of 22q11DS participants were taking medications at the time of the scan, and thus it is unlikely that medication effects played a role in the observed findings. Another concern, present across rs-fcMRI studies in clinical populations, relates to head movement. We movement-scrubbed all data and used movement (% frames scrubbed) as a covariate across all analysis, which did not alter the observed findings. Furthermore, motion parameters did not significantly differ between 22q11DS and typically developing control participants (**Table 1**) and rs-fcMRI effects were not related to head movement or signal-to-noise ratio (SNR) (**Table 2**). Finally, we studied subjects who, by virtue of a highly penetrant CNV, were at elevated risk for psychosis (and other neuropsychiatric symptoms). Given the young age of many of the study participants, current findings cannot address the question of whether the magnitude of thalamic-hippocampal dysconnectivity is indeed associated with subsequent risk for the development of psychosis. Importantly, the classification results indicate robust sensitivity-specificity, which may aid such prediction. Prospective longitudinal studies are currently underway to address this key knowledge gap.

### Conclusions

This study leverages the well-characterized genetic etiology of 22q11DS, thus providing a robust high-penetrance model to guide and test mechanistic hypotheses regarding disrupted brain development and subsequent consequences for circuit dysfunction leading to neuropsychiatric symptoms. Our findings offer the first evidence for reciprocal disruption of thalamic and hippocampal functional connectivity with cortical regions in this genetic risk model. Notably, the observed findings pinpoint an anterior axis of thalamic-hippocampal systems in line with animal model observations, which yield a robust classifier that can be refined for longitudinal risk prediction. These findings suggest that ongoing focus on thalamic-hippocampal circuit interactions in 22q11DS patients and in animal models can guide translational development of targeted and mechanistically informed neural markers and subsequent therapeutics.

## Acknowledgements

Aspects of the data were provided by the Human Connectome Project, WU-Minn Consortium (Principal Investigators: David Van Essen and Kamil Ugurbil; 1U54MH091657) funded by the 16 NIH Institutes and Centers that support the NIH Blueprint for Neuroscience Research; and by the McDonnell Center for Systems Neuroscience at Washington University. This work was supported by the NIH via awards DP5-0D012109 (Anticevic), R01-MH108590 (Anticevic), R01-MH112189 (Anticevic), R01 MH085953 (Bearden), U54 EB020403 (Bearden), T32MH073526 (Lin), as well as the Brain and Behavior Foundation (NARSAD) Independent Investigator grant (Anticevic) and the Joanne and George Miller Family Endowed Term Chair (Bearden).

